# ClpP-dependent protein quality control supports antibiotic persistence in *Campylobacter jejuni* through bioenergetic homeostasis

**DOI:** 10.1101/2024.07.15.603561

**Authors:** Jinsong Feng, Shenmiao Li, Yaxi Hu, Martin Stahl, Lina Ma, Katelyn Knuff-Janzen, Kaidi Wang, Yihan He, Bruce A. Vallance, Michael E. Konkel, B. Brett Finlay, Xiaonan Lu

**Affiliations:** Food, Nutrition, and Health Program, Faculty of Land and Food Systems, The University of British Columbia, Vancouver, British Columbia, V6T 1Z4, Canada; Michael Smith Laboratories, The University of British Columbia, Vancouver, British Columbia, V6T 1Z4, Canada; Department of Food Science and Nutrition, Zhejiang University, Hangzhou, Zhejiang, 310058, China; Department of Food Science and Agricultural Chemistry, Faculty of Agricultural and Environmental Sciences, McGill University, Sainte-Anne-de-Bellevue, Quebec, H9X 3V9, Canada; British Columbia Children’s Hospital Research Institute, Vancouver, British Columbia, V6H 3V4, Canada; School of Molecular Biosciences, College of Veterinary Medicine, Washington State University, Pullman, WA 99164, USA

## Abstract

Bacterial persistence enables survival during lethal antibiotic exposure and is implicated in recurrent infections, yet the physiology underlying bacterial persistence in many pathogens remains poorly defined. Here we show that exposure of *Campylobacter jejuni* to ampicillin or ciprofloxacin generates an antibiotic-persistent subpopulation. Rather than undergoing global metabolic shutdown, persister cells adopted a metabolically constrained state characterized by selective maintenance of oxidative phosphorylation and bioenergetic metabolism through coordinated proteostasis control. The ATP-dependent protease ClpP was essential for entry into this persistent state. Loss of ClpP disrupted proteostasis of the electron transport chain, specifically impaired *bd*-like terminal oxidase integrity, and reduced survival *in vivo* and in macrophages. These findings identify ClpP-dependent maintenance of redox and bioenergetic homeostasis as critical determinants of *C. jejuni* persistence and highlight metabolic remodeling as a defining feature of antibiotic tolerance. These insights may inform future therapeutic strategies aimed at disrupting persistence and improving antibiotic efficacy.

## INTRODUCTION

Since Bigger’s original observations, antibiotic persistence has been recognized as a phenotypic and non-heritable survival strategy by which a small subpopulation of bacteria survives otherwise lethal antibiotic exposure; these cells are termed persisters [1, 2]. Persisters have since been documented across diverse bacterial species and increasingly associated with the recalcitrance and relapse of bacterial infections [3–5]. Persistence is now recognized as a manifestation of phenotypic heterogeneity associated with variations in growth rate and adaptive physiological states, rather than a genetically encoded resistance phenotype [6–9].

Accumulating evidence suggests that bacterial metabolic state plays a central role in antibiotic tolerance. Bioenergetic and redox homeostasis underpin respiration, energy conservation, and metabolic processes required for growth and stress adaptation [10–12]. Bactericidal antibiotics disrupt essential cellular functions and often trigger energy-demanding compensatory responses, while numerous studies indicate that metabolic state both reflects and influences antibiotic lethality and survival [12–16]. Persister formation has frequently been associated with slowed growth, reduced biosynthetic activity, and low-energy states, but these features are increasingly viewed as part of a continuum of adaptive physiological states, rather than a binary switch between active and dormant cells [2, 17–19]. ATP depletion has been linked to increased survival during antibiotic challenge in *Staphylococcus aureus* and *Escherichia coli*, and their metabolic state can predict antibiotic lethality more accurately than growth rate, when growth and metabolism are experimentally uncoupled [8, 18–20].

Metabolic perturbation is also closely connected to protein homeostasis. ATP depletion has been linked to protein aggregation and persistence-associated dormancy [21, 22]. In bacteria, protein aggregation arises from proteostatic stress and has been associated with altered growth, loss of protein function, stress tolerance, and pathogenicity [22–25]. Deletion of *pspA* in stationary-phase *E. coli* lowers intracellular ATP levels, accelerates protein aggregation, and increases persistence, whereas glucose supplementation partially restores ATP levels and reduces persister formation [26]. ATP-dependent protein aggregation has also been shown to correlate with dormancy depth and lag time, while the DnaK–ClpB disaggregation machinery is required for efficient resuscitation [22]. These findings suggest that protein aggregation may link bioenergetic stress, slowed growth, dormancy depth, and antibiotic survival.

Proteostasis has also been implicated in toxin–antitoxin (TA) models of persistence, in which antitoxin degradation and toxin activation were proposed to promote reversible growth arrest [27]. In *E. coli*, MazF-mediated persistence was reported to require the ClpP and Lon proteases [28]. However, the broader contribution of type II TA systems to antibiotic persistence remains debated, as deletion of ten chromosomal type II TA loci in *E. coli* did not impair persistence to ampicillin or ofloxacin [29]. Still, individual toxins can influence persistence-associated physiology. For example, the SOS-regulated type I membrane toxin TisB promoted dormancy- linked changes, including dissipation of the proton motive force, loss of pH homeostasis, and in some contexts, protein aggregation that prolongs dormancy [30, 31].

In parallel, studies of bactericidal antibiotic lethality proposed a shared oxidative death pathway mediated by reactive oxygen species (ROS) [16]. Although influential, this model has been recently revised, as subsequent work demonstrated that bactericidal activity can occur independently of ROS and under oxygen-limited conditions [32, 33]. The field now recognizes that antibiotic stress perturbs respiration, electron flow, and cellular redox physiology in a context-dependent manner, with ROS representing one possible downstream consequence rather than a universal mechanism [8, 34–36]. From this perspective, redox imbalance provides a more robust lens through which to understand how antibiotic stress reshapes bacterial physiology and survival.

These questions are particularly relevant in *Campylobacter jejuni*, a leading cause of foodborne bacterial gastroenteritis worldwide. Compared with many bacterial pathogens, *C. jejuni* has a relatively small genome (∼1.6–1.7 Mb) and lacks several classical stress-response regulators, including RpoS, OxyR, and SoxRS [37, 38]. Despite these apparent limitations, *C. jejuni* survives diverse environmental and host-associated stresses and can form antibiotic-tolerant persisters. A clinical case study described relapsing infections associated with long-term colonization despite antibiotic treatment [4], and additional studies showed that exposure to antibiotics, such as ciprofloxacin, penicillin G, or tetracycline can generate tolerant or persister-like subpopulations [39–41]. In penicillin G-treated cells, Morcrette and co-authors identified a subpopulation of *C. jejuni* with increased fluorescence after staining with a redox-sensitive dye, suggesting altered redox activity or respiratory remodeling [40]. Nevertheless, the physiological mechanisms underlying the persistence in *C. jejuni* remain poorly defined.

Here, we investigated how metabolic adaptation supports antibiotic persistence in *C. jejuni*, with a particular focus on the coupling between redox homeostasis, bioenergetic remodeling, and proteostasis. Using antibiotics with strong metabolic dependencies, together with genetic, physiological, and metabolic analyses, we investigated whether persistence reflects generalized metabolic suppression or instead a more selective remodeling of survival-associated pathways. Our findings identify ClpP-dependent proteostasis as a central determinant of antibiotic tolerance and suggest that antibiotic-tolerant *C. jejuni* undergoes respiratory and redox remodeling rather than a uniform metabolic shutdown.

## RESULTS

### *C. jejuni* forms an antibiotic-tolerant subpopulation with bactericidal antibiotic treatment

To determine whether *C. jejuni* can develop antibiotic tolerance when exposed to bactericidal antibiotics with distinct primary targets, we performed time-killing assays using ampicillin and ciprofloxacin in four representative *C. jejuni* strains (F38011, 81-116, 87-95, 11168) with slightly different genetic backgrounds. Under antibiotic concentrations that were bactericidal, we observed marked heterogeneity within the *C. jejuni* populations. Although most cells were rapidly eliminated, a small surviving fraction exhibited a substantially slower killing rate, resulting in a biphasic killing pattern (**Fig. 1a, b**). This biphasic pattern was consistently observed across all four strains tested, indicating that persister formation was not limited to a single strain background in our strain set. To exclude the possibility that the surviving cells represented transiently resistant variants, we repeatedly challenged the survivors with the same antibiotics over four successive rounds. No increase in MIC was detected, indicating that the surviving subpopulation had not acquired heritable resistance (**Extended Data Figure 1** and **Supplementary Table 1**). We also ruled out the possibility that reduced killing at later time points resulted from antibiotic degradation by pre-incubating the antibiotics under the same conditions used in the killing assays for 6 h prior to bacterial exposure. This pre-incubation did not alter the MICs of either antibiotic. Collectively, these results demonstrated that *C. jejuni* transiently formed an antibiotic-tolerant subpopulation under bactericidal antibiotic stress, and that the survival of this subpopulation reflected physiological adaptation, rather than stable resistance mutations.

**Figure 1.**
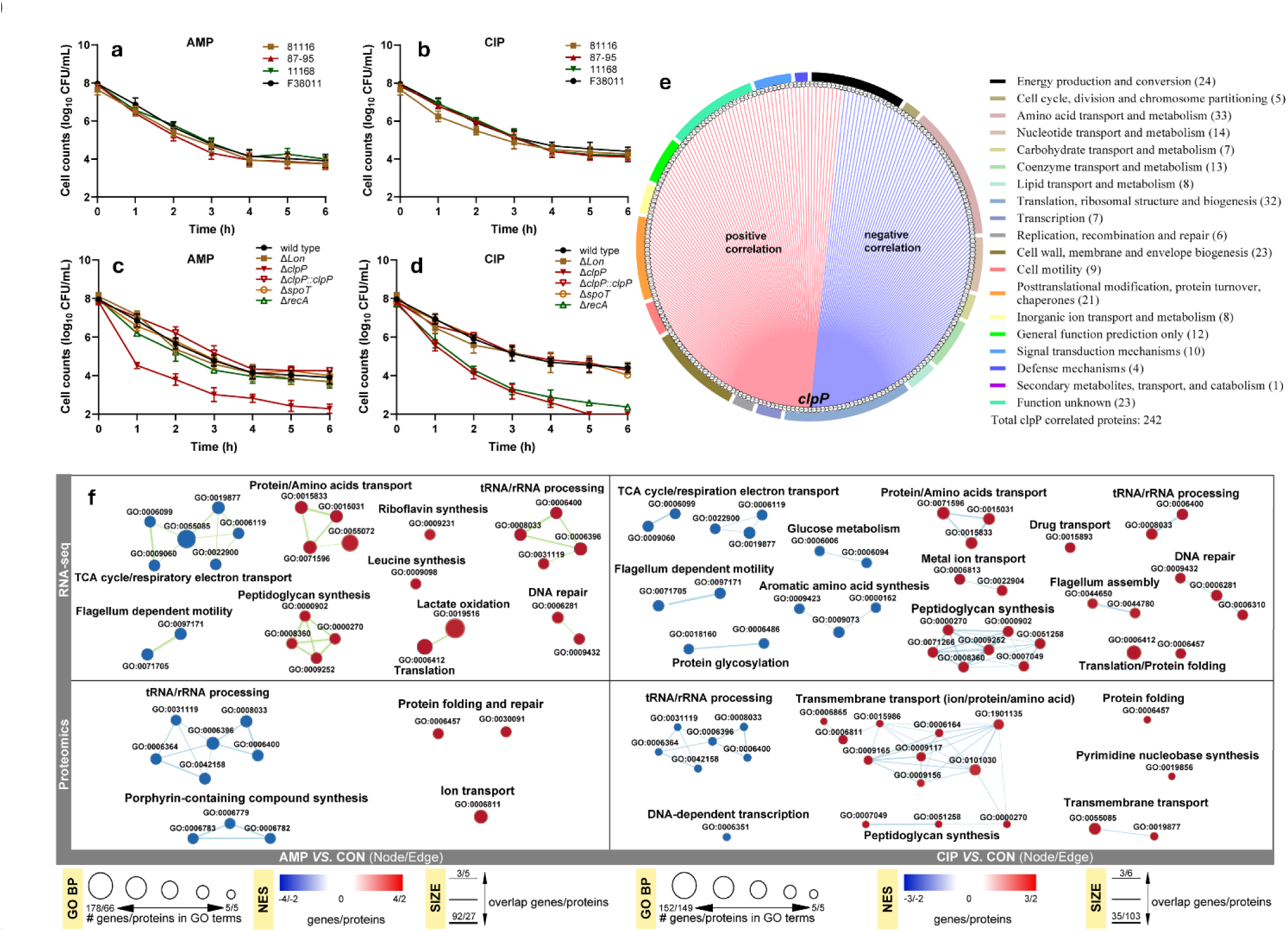
*clpP* is critical for *Campylobacter jejuni* persisters to tolerate ampicillin (AMP) and ciprofloxacin (CIP). **(a, b)** Time-killing curves of AMP (a) and CIP (b) against four *C. jejuni* strains: 81116 (dark yellow squares), 87-95 (red triangles), 11168 (green inverted triangles), and F38011 (black circles). Cultures were grown to an optical density of 0.3 and then diluted to approximately log 8 CFU/mL in Mueller–Hinton (MH) broth containing either ampicillin (100 μg/mL) or ciprofloxacin (1 μg/mL). **(c, d)** Time-killing curves of AMP (c) and CIP (d) against wild-type F38011 (black circles), the *Δlon* mutant (dark yellow squares), *ΔclpP* mutant (red inverted triangles), *ΔspoT* mutant (orange open circles), *ΔrecA* mutant (green open triangles), and the *clpP*-complemented strain (*ΔclpP::clpP*; red open inverted triangles). **(e)** Cluster of Orthologous Groups (COG) classification of proteins strongly correlated with *clpP*. A total of 241proteins were identified based on an absolute Pearson correlation coefficient >0.8 and *p* < 0.05. COG analysis grouped these *clpP*-correlated proteins into 19 functional categories; the number of proteins assigned to each category is indicated in parentheses. **(f)** Gene set enrichment analysis (GSEA) of transcriptomic and proteomic profiles of *C. jejuni* persisters. GSEA was performed using pre-ranked lists of all genes or proteins based on differential expression between antibiotic-selected persister cells and untreated cells. The cumulative distribution function was generated using 1,000 random gene set membership permutations. Significantly enriched terms from transcriptomic and proteomic analyses were visualized in Cytoscape 3.7.1. GO enrichments were constructed using a stringent pathway similarity threshold (combined Jaccard and overlap coefficient = 0.6) and manually curated to highlight representative biological process (GO BP). Node size corresponds to gene set size, and node color indicates enrichment in antibiotic-selected persister cells (red; upregulated after antibiotic treatment) or untreated cells (blue; downregulated after antibiotic treatment). Edge thickness represents the degree of gene overlap between connected gene sets. Data are presented as mean ± SD from at least three independently grown cultures.

### Multi-omics profiling reveals proteostasis and respiratory remodeling in tolerant cells

To characterize the physiological state of antibiotic-tolerant *C. jejuni* cells, we performed transcriptomic and proteomic analyses on the tolerant fraction collected after ampicillin or ciprofloxacin treatment. Gene set enrichment analysis of the transcriptomic data revealed consistent enrichment of pathways associated with protein catabolism and quality control, SOS and stringent stress responses, peptidoglycan synthesis, and RNA processing. In contrast, pathways related to flagellum-dependent motility, the TCA cycle, and respiration/electron transport were under-represented (**Fig. 1f**). These results indicate that antibiotic-tolerant cells do not simply undergo global metabolic shutdown, instead they adopted a selective regulatory program characterized by enhanced stress adaptation and altered respiratory activity. Proteomic profiling further supported this interpretation, indicating preservation of stress and homeostasis-associated functions despite broad metabolic restriction. Although transcriptomic and proteomic changes were not fully concordant, several key functional trends were consistent across both datasets. Notably, pathways involved in protein folding and quality control remained enriched under both antibiotic treatments, suggesting sustained investment in proteostasis despite the energetically restricted state of tolerant cells. Ciprofloxacin exposure induced broader proteomic changes than ampicillin, yet both treatments converged on a shared signature involving protein homeostasis and metabolic reprogramming.

### ClpP is critical for antibiotic tolerance in *C. jejuni*

To investigate the pathways involved in antibiotic tolerance, we generated a panel of *C. jejuni* mutants on the F38011 background, including mutants defective in the stringent response (Δ*spoT*) [42], SOS response (Δ*recA*) [43], and ATP-dependent proteolysis (Δ*lon* and Δ*clpP*) [44]. We then compared their survival under ampicillin and ciprofloxacin treatment (**Fig. 1c, d**).

Primers and plasmids used for strain construction are listed in **Supplementary Table 2**. Deletion of *spoT* or *recA* did not affect tolerance to ampicillin, whereas deletion of *recA* selectively reduced survival under ciprofloxacin treatment, consistent with the established role of RecA in DNA damage repair. In contrast, deletion of *clpP* markedly compromised survival under both ampicillin and ciprofloxacin treatment, while deletion of *lon* had no significant effect under either antibiotic condition. Importantly, the Δ*clpP* mutant displayed MIC values comparable to those of the wild-type strain and showed no significant growth defect under non-stressed conditions. Therefore, the reduced survival of the Δ*clpP* mutant cannot be attributed to altered intrinsic susceptibility or impaired growth but instead reflects a specific defect in antibiotic tolerance. Collectively, these results identify ClpP as a key determinant of antibiotic tolerance in *C. jejuni*.

### *clpP*-associated protein regulation links tolerance to respiratory homeostasis

To further refine the *clpP*-associated regulatory module, we stratified proteins according to their abundance correlation with *clpP* and their predicted disorder tendency. The *clpP*-centered protein co-expression analysis identified 241 proteins whose abundance was strongly correlated with *clpP* expression (absolute Pearson correlation coefficient > 0.8, *p* < 0.05; **Supplementary Table 3**). These proteins were predominantly associated with pathways related to 1) energy production and conversion, 2) amino acid transport and metabolism, 3) translation, ribosomal structure, and biogenesis, 4) cell wall, membrane and envelope biogenesis, and 5) post-translational modification, protein turnover, and chaperones (**Fig. 1e**). These *clpP*-correlated proteins were overall better maintained during antibiotic treatment than non-correlated proteins, showing higher relative abundance under both ampicillin and ciprofloxacin exposure (**Fig. 2a, b**). We then analyzed the disorder tendency of the *clpP*-correlated proteins using DFLpred [45] and classified them into high-, medium-, and low-disorder groups. This stratification was used to determine whether the ClpP-associated module was preferentially linked to proteins with greater predicted structural flexibility, given that proteins with elevated aggregation risk are often subject to tighter transcriptional, translational and degradation control as part of cellular proteostasis strategies [46]. Among the *clpP*-correlated proteins detected in both conditions, 83 were classified as high-disorder, 135 as medium-disorder, and 22 as low-disorder (**Fig. 2c, d**; **Table S3**). In contrast, among the 132 low-disorder proteins detected, only 22 were *clpP*-correlated whereas 110 were non-correlated, indicating that ClpP-associated regulation was preferentially linked to proteins with higher disorder tendency. Consistently, the high-disorder subset showed the highest relative abundance within the *clpP*-correlated cluster under both antibiotic treatments (**Fig. 2c, d**). Comparison of mRNA and protein fold changes for the *clpP*-correlated/high-disorder subset further revealed an asymmetric transcript–protein relationship, in which positive mRNA changes were more closely reflected at the protein level than negative mRNA changes (**Fig. 2e, f**). Mapping these features onto KEGG pathways further showed that the *clpP*-correlated/high-disorder module was associated with selectively maintained functions in glycolysis/gluconeogenesis, pyruvate metabolism, the TCA cycle, oxidative phosphorylation, and amino acid metabolism. In contrast, functions related to fatty acid elongation, biotin biosynthesis, and parts of histidine, lysine, and glutamate metabolism were suppressed (**Fig. 2g**).

**Figure 2.**
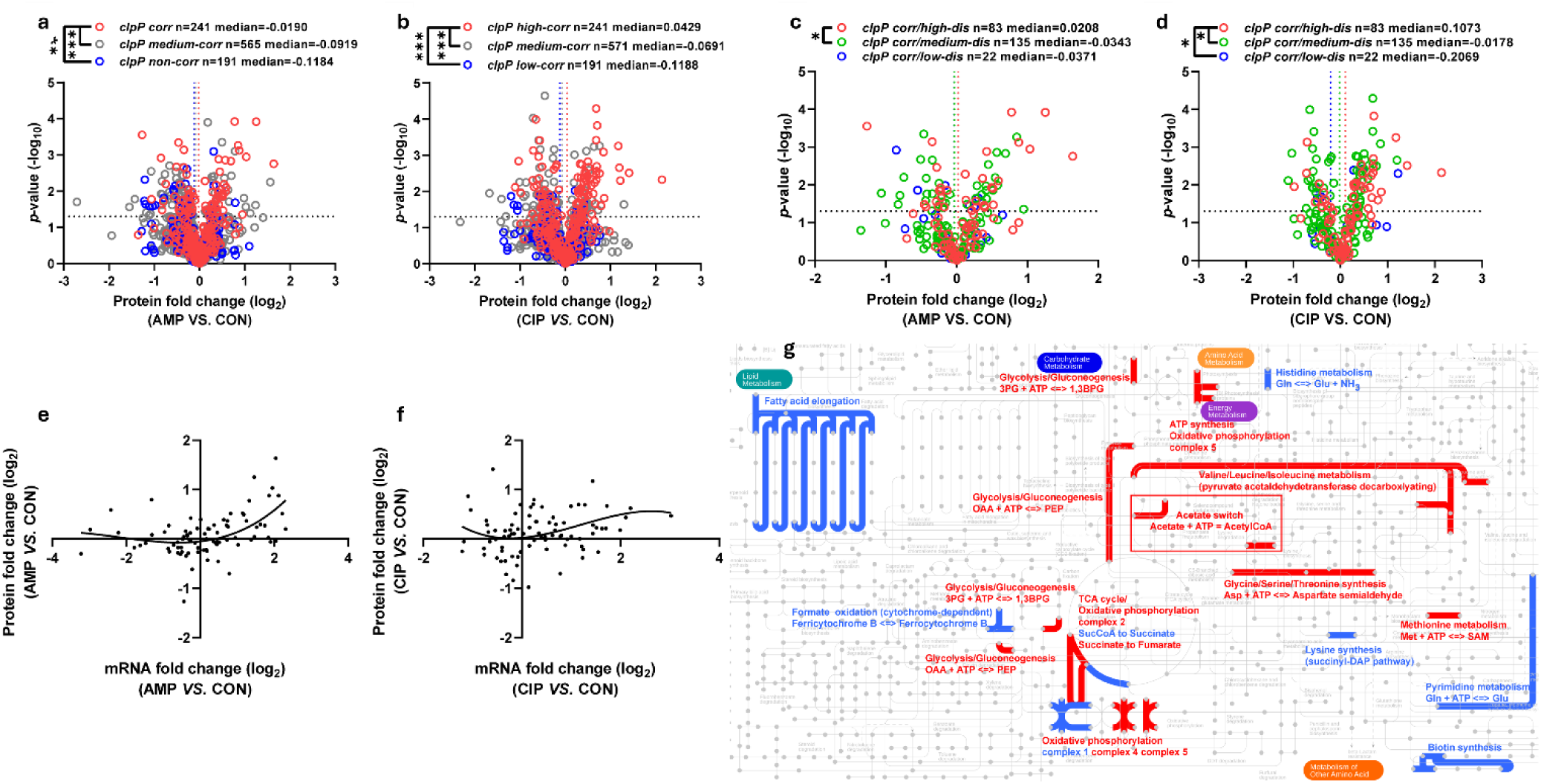
*clpP*-associated high-disorder proteins define metabolically active functions in *C. jejuni* persisters. **(a, b)** Distribution of protein fold changes in *C. jejuni* F38011 persisters after ampicillin (a) or ciprofloxacin (b) treatment, stratified by the correlation between protein abundance and *clpP* expression. Pearson correlation analysis was used to determine abundance correlation with *clpP*. *clpP*-correlated features were defined as proteins with an absolute correlation coefficient ∣ *r* ∣> 0.8 and *p* < 0.05, whereas non-correlated features were defined as proteins with ∣ *r* ∣< 0.2and *p* < 0.05. **(c, d)** Distribution of protein fold changes among *clpP*-correlated proteins was further stratified by predicted disorder tendency after ampicillin (c) or ciprofloxacin (d) treatment. Disorder tendency was predicted using DFLpred based on the proportion of disordered flexible linker residues within each protein sequence. High disorder proteins were determined as disordered flexible linker residues accounting for >30% of the total sequences. Non-disorder proteins were determined as disordered flexible linker residues accounting for <10% of the total sequences. Among the *clpP*-correlated proteins detected in both conditions, 83 were classified as high-disorder, 135 as medium-disorder, and 22 as low-disorder. **(e, f)** Relationship between mRNA and protein log2 fold changes for *clpP*-correlated/high-disorder features in persisters after ampicillin (e) or ciprofloxacin (f) treatment. Differentially expressed genes and proteins were compared across the corresponding pairwise contrasts. **(g)** Global metabolic map derived from *clpP*-correlated/high-disorder features. Red indicates promoted activity, whereas blue indicates inhibited activity.

### ClpP supports the antibiotic-induced respiratory response and *bd*-like terminal oxidase integrity

Because multi-omics analysis pointed to substantial remodeling of respiration- and electron transport-associated pathways, we next assessed respiratory activity by measuring the oxygen consumption rate (OCR) using Seahorse analysis. Antibiotic exposure triggered a transient increase in oxygen consumption, manifested as a distinct OCR peak (**Fig. 3a**). Notably, the wild-type F38011 strain exhibited a stronger respiratory response than the Δ*clpP* mutant and maintained higher overall respiratory activity throughout the post-treatment period. Normalization of OCR to viable cell counts showed that antibiotic-treated wild-type cells still exhibited higher respiratory activity than the Δ*clpP* mutant (**Fig. 3b**). Thus, the reduced OCR observed in Δ*clpP* cannot be attributed solely to differences in viable cell number after treatment. Rather, these data support a role for ClpP in maintaining bioenergetic homeostasis during antibiotic stress.

**Figure 3.**
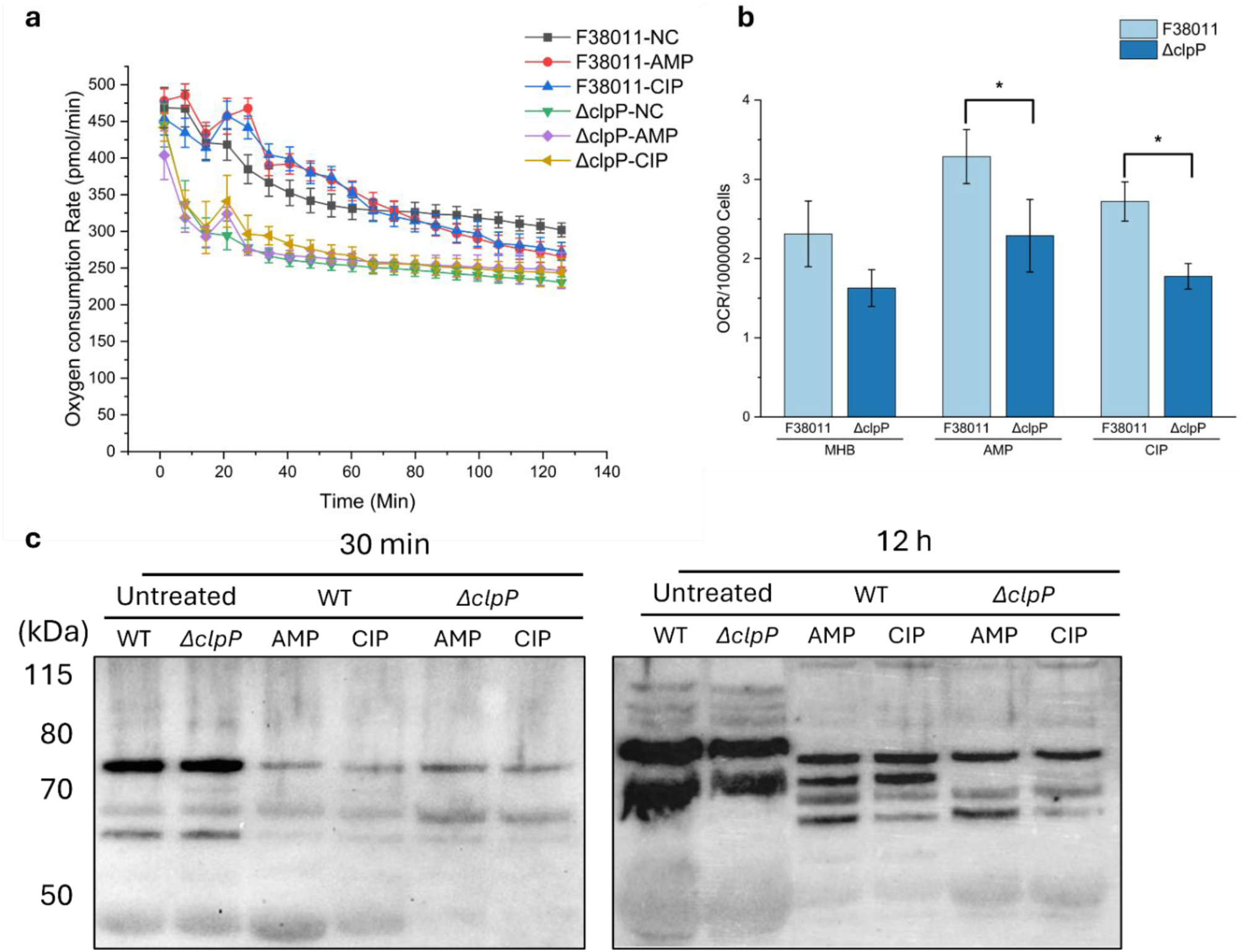
ClpP supports the antibiotic-induced respiratory response and bd-like terminal oxidase integrity in *C. jejuni*. **(a)** Real-time changes in oxygen consumption rate (OCR; pmol O_2_/min) in wild-type F38011 and the *ΔclpP* mutant in response to antibiotic treatment were measured using a Seahorse XFe Extracellular Flux Analyzer. OCR was monitored over time in untreated control cells (NC) and in cells exposed to ampicillin (AMP) or ciprofloxacin (CIP) at 20 μg/mL and 0.2 μg/mL, respectively. Antibiotic treatment induced a transient increase in respiration, and the wild-type strain displayed a stronger and more sustained OCR response than the *ΔclpP* mutant. Data represent mean ± SD of 12 technical replicates. **(b)** Normalized OCR per live cell. (\**p* < 0.05). **(c)** Western blot analysis of the bd-like terminal oxidase subunit CydA in wild-type F38011 and *ΔclpP* cells exposed to ampicillin or ciprofloxacin at 10 μg/mL and 0.1 μg/mL for 30 min or 12 h. Untreated cells served as no-antibiotic controls. Lane order in each panel is: untreated WT, untreated *ΔclpP*, WT + AMP, WT + CIP, *ΔclpP* + AMP, and *ΔclpP* + CIP. At 30 min, CydA-associated signals were weak across antibiotic-treated samples. By 12 h, wild-type cells retained a clearer multi-band CydA pattern, whereas *ΔclpP* cells showed an altered banding pattern characterized by reduction of the lower CydA-associated bands despite retention of the upper band. Molecular mass markers (kDa) are indicated on the left.

*C. jejuni* encodes two terminal oxidases, the high-affinity *cbb_3_*-type oxidase CcoNOQP and a cyanide-insensitive *bd*-like oxidase encoded by *cydAB* (also referred to as *cioAB*) [47, 48]. Previous studies indicate that the *cbb*_3_ branch is the major high-affinity oxidase supporting microaerobic respiration, whereas the *bd*-like branch is a lower-affinity oxidase whose expression increases under higher oxygen availability and is more commonly associated with stress-adaptive respiration [48, 49]. Transcriptomic analysis further showed selective remodeling of central metabolic and respiratory functions in antibiotic-treated cells. Selected TCA cycle genes were downregulated, and the downregulated respiratory genes were primarily enriched in oxidative phosphorylation and electron transport chain complex I, II, III, and IV (CcoNOQP). By contrast, genes associated with ATP synthase complex V were upregulated (**Extended Data Figure 2**). In addition, *cydAB* (*cj0081* and *cj0082*) was upregulated in both antibiotic-treated populations (**Table S6, Extended Data Figure 2**). These patterns suggest altered respiratory coupling, reduced proton-translocating respiratory capacity and sustained energetic demand.

We next examined the *bd*-like terminal oxidase subunit CydA by Western blotting following exposure to antibiotics at MIC. Untreated controls displayed the characteristic CydA banding pattern under unstressed conditions. At 30 min, CydA-associated signals were weak across antibiotic-treated samples, limiting interpretation at this early time point (**Fig. 3c, left**). By 12 h, a clearer difference emerged between strains. Specifically, wild-type cells displayed a reproducible multi-band CydA pattern spanning approximately 65-75 kDa, including a major upper band at ∼75 kDa and several lower-molecular-weight bands between ∼60 and 70 kDa. In contrast, the Δ*clpP* mutant showed an altered pattern characterized by loss of the band at ∼70 kDa, while retaining the major upper band (**Fig. 3c, right**). Because CydA is a membrane-embedded subunit of the *bd*-like terminal oxidase, detection of multiple CydA-associated bands is not unexpected, although the basis of these species remains unresolved. This shift in the CydA banding pattern suggests that the respiratory defect in the Δ*clpP* mutant involves altered CydA-associated terminal oxidase integrity rather than complete loss of the *bd*-like terminal oxidase.

### Antibiotic-tolerant cells undergo redox-linked metabolic and membrane lipid remodeling

To further define the metabolic state of antibiotic-tolerant cells, we performed metabolomic and lipidomic analyses to examine changes in both hydrophilic and hydrophobic metabolites (**Fig. 4a, d**, **Tables S4 and S5**). The metabolomic data collectively revealed an overall decrease in the abundance of metabolites associated with core metabolic pathways. Consistent with the regulatory patterns inferred from transcriptomics and proteomics, metabolites involved in biotin biosynthesis and histidine, lysine, and glutamate metabolism were significantly downregulated (**Fig. 4b, c** and **Figure S3**). Importantly, key metabolites involved in glycolysis, the TCA cycle, and amino acid metabolism were also overtly depleted, indicating broad suppression of central carbon metabolism. In contrast, several metabolites involved in nucleotide sugar metabolism and the pentose phosphate pathway, including UDP-xylose, UDP-D-glucose, and UDP-glucuronate, accumulated under both ampicillin and ciprofloxacin treatment. To further interpret this shift, we constructed a core metabolic map of *C. jejuni* (**Fig. 5a**). This analysis suggested that the apparent upregulation of glycolysis/gluconeogenesis-related reactions did not necessarily reflect increased glycolytic flux, but rather a redirection of carbon toward nucleotide sugar metabolism. The tolerant cells also displayed a shunted TCA cycle, depletion of the glutamate–glutamine cycle, and accumulation of redox-associated metabolites such as S-adenosylmethionine, riboflavin-5-phosphate, and menaquinone-6. Together, these findings support a metabolically constrained but redox-oriented tolerant state.

**Figure 4.**
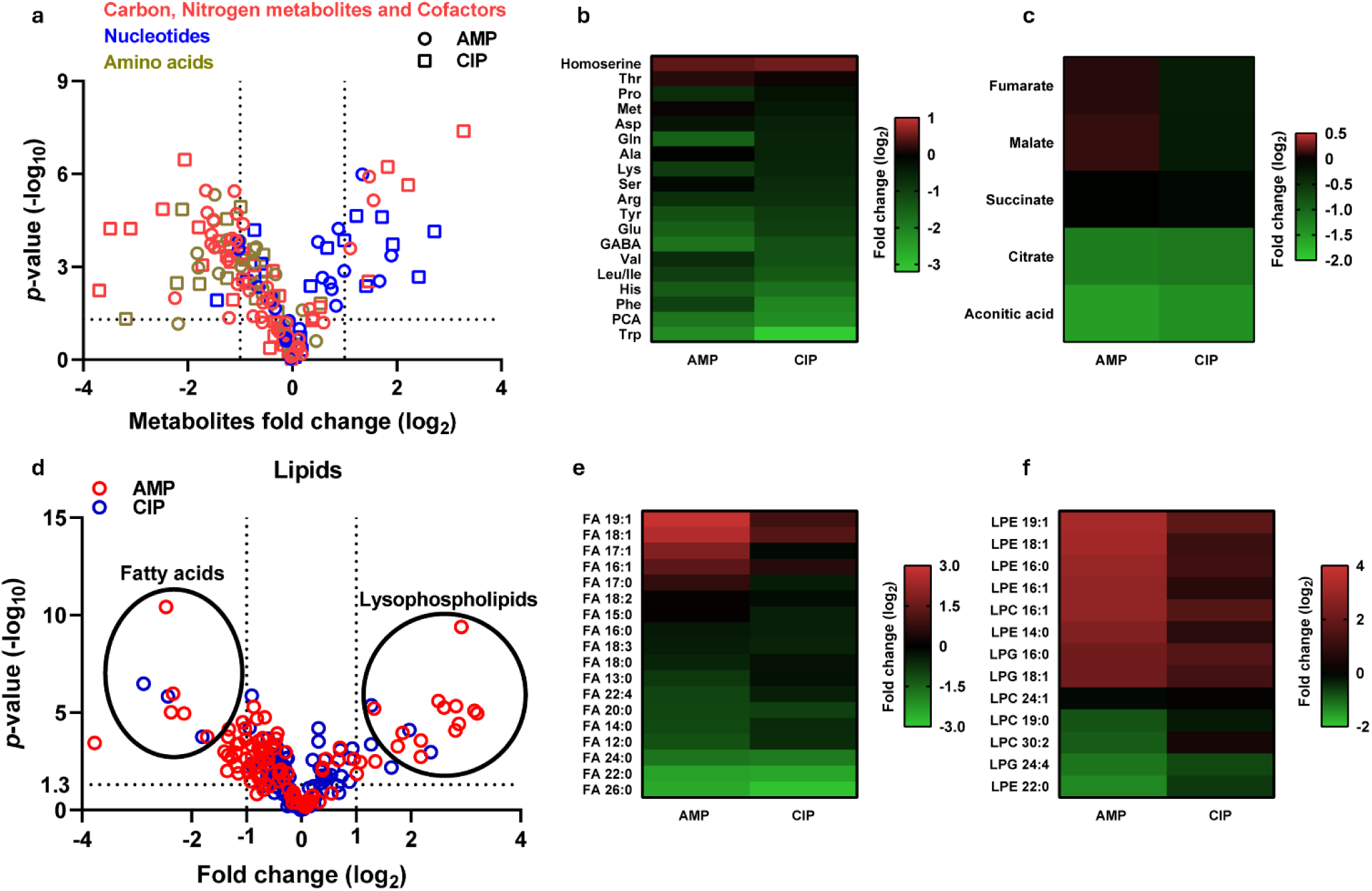
*C. jejuni* F38011 persisters exhibited distinct metabolomic and lipidomic features compared with untreated growing cells. **(a)** Volcano plot showing changes in relative metabolite abundance in persister cells selected by AMP or CIP compared with untreated cells. Metabolites are color-coded by functional class, including carbon, nitrogen, and cofactor-related metabolites, nucleotides, and amino acids. **(b)** Heatmap showing relative abundance changes of selected amino acids in AMP and CIP selected persister cells. **(c)** Heatmap showing relative abundance changes of selected TCA cycle intermediates in AMP and CIP selected persister cells. **(d)** Volcano plot showing changes in relative lipid abundance in persister cells selected by AMP or CIP compared with untreated cells. Fatty acids and lysophospholipids are highlighted. **(e)** Heatmap showing relative abundance changes of fatty acids in *C. jejuni* persister cells. **(f)** Heatmap showing relative abundance changes of lysophospholipids in *C. jejuni* persister cells.

**Figure 5.**
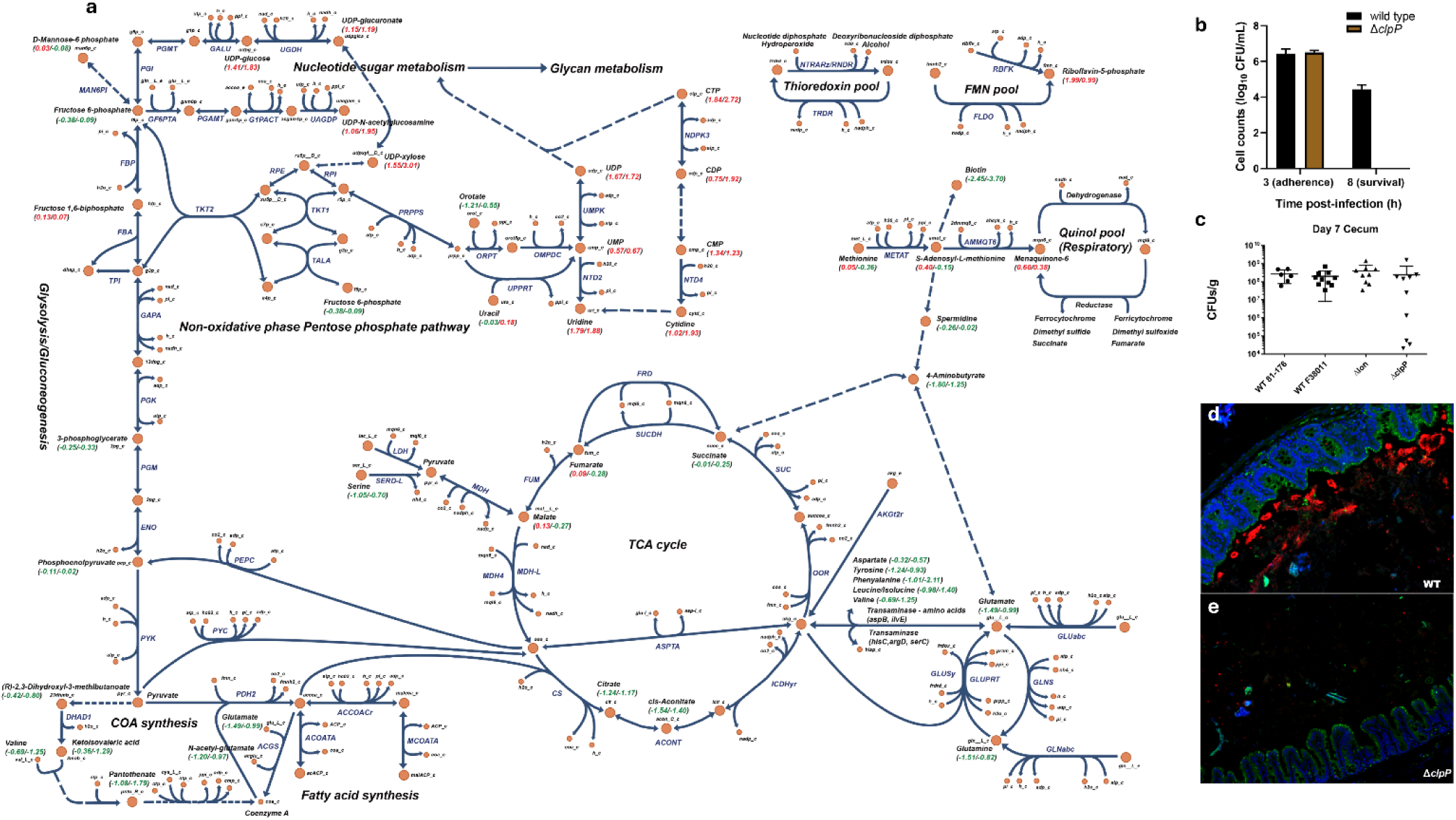
Metabolic reprogramming of *C. jejuni* persisters and reduced host-associated survival of the *clpP* mutant. **(a)** Core metabolic map illustrating metabolic reprogramming in *C. jejuni* persister cells relative to untreated growing cells. Relative metabolite abundance changes are annotated on the network, with red values indicating increased abundance (log2 fold change) and green values indicating decreased abundance (log fold change). Left: AMP; Right: CIP. **(b)** Survival of wild-type F38011 and the persistence-compromised *clpP* mutant in macrophages. **(c)** Cecal colonization levels of wild-type F38011 and the *clpP* mutant in *sigirr*^-/-mice. **(d, e)** Representative images showing localization of wild-type F38011 (d) and the *clpP* mutant (e) in the cecum of *sigirr*^-/- mice.

Lipidomic analysis further revealed pronounced membrane remodeling in antibiotic-tolerant cells. Fatty acids with chain lengths predominantly ranging from C_14_ to C_18_ accumulated under both antibiotic treatments (**Fig. 4e, f**), whereas biotin, a cofactor required for fatty acid biosynthesis, was significantly depleted (**Figure S3**), suggesting that fatty acid accumulation primarily resulted from catabolic processes rather than de novo synthesis. In addition, a broad range of lysophospholipids, particularly species with C_16_ and C_18_ chains, was significantly enriched in tolerant cells. This was accompanied by upregulation of *pldA* (**Table S7**), which encodes an outer membrane phospholipase that can hydrolyze oxidized phospholipids to generate lysophospholipids and free fatty acids under stress conditions [50, 51]. Consistent with this, oxidized phosphatidylethanolamine intermediates were also detected (**Figures S6** and **S7**), suggesting that antibiotic treatment promoted phospholipid oxidation and lipid turnover even though bulk lipid peroxidation was not markedly increased (**Figures S4** and **S5**). These lipid changes indicated that the tolerant state is accompanied by membrane turnover and phospholipid remodeling.

### Loss of ClpP compromises redox homeostasis and host survival

Given the central role of ClpP in antibiotic tolerance and the extensive redox- and membrane-associated remodeling observed in tolerant cells, we next examined whether loss of ClpP disrupts homeostasis more broadly. Deletion of *clpP* increased sensitivity to oxidative stress and elevated lipid peroxidation, indicating that ClpP contributes to the maintenance of both redox balance and membrane integrity (**Fig. 4 and Figure S4**). This phenotype is consistent with a role for ClpP-dependent protein quality control in limiting damage accumulation under stress conditions, and in preserving processes linked to respiratory function. We then assessed whether this defect affected host-associated survival. In RAW 264.7 macrophages, the wild-type and Δ*clpP* mutant strains exhibited similar adherence at 3 h post-infection, but the Δ*clpP* mutant rapidly lost viability thereafter and became undetectable by 8 h post-infection (**Fig. 5b**). To further evaluate the pathogenic relevance of this phenotype *in vivo*, we infected Sigirr-deficient mice, which were previously shown to be susceptible to enteric *C. jejuni* infection [52], and measured intestinal pathogen burdens. As expected, Sigirr-deficient mice were heavily colonized by wildtype *C. jejuni* F38011. Although the Δ*clpP* mutant retained some colonization capacity, its bacterial load was significantly lower than that of the wild-type strain (**Fig. 5c**). Histological analysis further showed that wild-type cells formed microcolonies on the epithelial surface, whereas the Δ*clpP* mutant was only rarely detected at the intestinal surface (**Fig. 5d, e**). Taken together with the *in vitro* stress phenotypes, these results extended the role of ClpP from antibiotic tolerance to survival in host-associated environments.

## DISCUSSION

Antibiotic tolerance has long been viewed as a stochastic survival phenomenon, but it is increasingly recognized as the outcome of coordinated physiological states that allow a subpopulation of bacterial cells to survive otherwise lethal antibiotic exposure [14, 53, 54]. This conceptual shift has also highlighted metabolism as a central determinant of antibiotic efficacy, tolerance, and the longer-term evolution of resistance [14, 53, 55]. Recent work further argues against viewing persistence simply as metabolic dormancy, instead emphasizing metabolically rewired and physiologically heterogeneous survival states [56]. Within this framework, our data support a view in which antibiotic tolerance in *C. jejuni* reflects active homeostatic remodeling rather than passive metabolic shutdown, with ClpP-dependent proteostasis emerging as a key feature of this state. This interpretation aligns with the broader role of ClpP in bacterial stress fitness, virulence, and proteome control across species, including *Listeria monocytogenes* and *Staphylococcus aureus* [57–60], and with mechanistic work showing that the ClpXP system can act on structurally diverse substrates through flexible recognition modes [61, 62]. It also aligns with growing interest in bacterial proteases as antimicrobial targets, including compounds that kill cells by dysregulating ClpP-mediated proteolysis [63].

Classical persistence-associated determinants include alarmone signaling, TA modules, stress-responsive proteases, and phenotypic switching networks [54, 64–66]. Our data suggest that antibiotic tolerance in *C. jejuni* depends less on a single canonical persistence pathway and more on the capacity of the cell to maintain physiological integrity during stress. Among the mutants tested, only Δ*clpP* showed compromised survival under both ampicillin and ciprofloxacin treatment, whereas Δ*recA* showed a condition-specific defect matching DNA damage repair, while Δ*spoT* and Δ*lon* did not phenocopy the survival defect. Because deletion of *clpP* did not alter MIC or basal growth, its phenotype is most consistent with a specific defect in tolerant-state survival. Tolerant cells showed enrichment of protein quality control and damage repair functions alongside repression of multiple growth-associated processes [10, 14, 65, 67, 68]. The partial transcript-protein discordance is mechanistically informative rather than problematic. Proteome allocation is not dictated solely by transcript abundance [69, 70], and translational regulation can further shape protein output even when transcript changes are incomplete or transient [71]. Recent work in *C. jejuni* has also revealed a more complex translatome and a broader role for translation-level regulation [72]. Proteostasis vulnerabilities are highly condition-specific [73], and proteases can support survival during growth arrest in a hierarchical manner, with ClpXP often playing a more dominant and nonredundant role than Lon [64]. Small-molecule studies likewise emphasize the importance of ClpP in tolerance, as dysregulation or hyperactivation of ClpP can eliminate persister populations [74, 75]. The enrichment of protein quality control across both datasets and the preferential association of ClpP with structurally vulnerable proteins support a model in which ClpP helps preserve a selective homeostatic proteome under proteotoxic stress [25, 76, 77]. Because repair and clearance of damaged proteins are energetically costly, this model also implies that tolerant cells remain selectively active rather than passively shut down. This view is reinforced by biochemical and single-molecule evidence that AAA+ protease systems engage substrates probabilistically and through diverse recognition modes [61, 62].

Functionally, this ClpP-associated module was enriched in central metabolism, oxidative phosphorylation, and amino acid metabolism, linking proteostasis to respiratory homeostasis. Rather than pointing to separate defects, the Seahorse and Western blot data converge on the view that antibiotic challenge imposed a respiratory-redox transition and that ClpP was required to maintain homeostasis through that transition [10, 78–82]. Given that *C. jejuni* physiology is tightly shaped by oxygen availability and a branched electron transport chain [47, 48, 83], and prior evidence has linked ClpXP to oxidative stress responses, redox sensing, and aerobic-anaerobic transitions [61], our findings support a role for ClpP in maintaining bioenergetic stability during antibiotic challenge.

The metabolomic and lipidomic data further define the tolerant state as one shaped by redox-linked metabolic adaptation and membrane remodeling. Depletion of central carbon metabolites together with accumulation of selected nucleotide-sugar intermediates and redox-associated metabolites points to redirected carbon use and a redox-oriented survival physiology, in line with evidence that pentose phosphate metabolism and redox-active metabolites can shape antibiotic lethality and tolerance [10, 84]. In parallel, accumulation of fatty acids and lysophospholipids, biotin depletion, *pldA* upregulation, and detection of oxidized phosphatidylethanolamine intermediates indicate membrane turnover and phospholipid remodeling. Because the membrane provides the structural platform for electron transport and proton translocation, these lipid changes are likely relevant to respiratory homeostasis [85]. They may also influence host interactions, given the emerging relevance of lysophosphatidylethanolamine in *C. jejuni* virulence [86]. Loss of ClpP also increased oxidative stress sensitivity and lipid peroxidation and reduced survival in macrophages and in the murine gut, indicating that ClpP-dependent functions that support antibiotic survival also contribute to fitness in host-associated environments, in line with prior work linking respiratory flexibility and niche adaptation in *Campylobacter* [47, 48, 83] and with evidence that ClpXP contributes to antioxidant control and population-level fitness maintenance in *S. aureus* [60].

Our data support a model in which antibiotic-tolerant *C. jejuni* cells exist in a metabolically constrained but selectively active state. We summarize this model in **Fig. 6**. Under basal microaerobic conditions, respiration supports efficient energy conservation while ClpP-dependent proteostasis maintains proteome integrity. Following bactericidal antibiotic exposure, surviving cells transition into a stress-associated tolerant state characterized by respiratory remodeling, redox-oriented metabolic adaptation, and increased reliance on protein quality control. In this state, ClpP-mediated clearance of damaged proteins helps preserve cellular homeostasis, and disruption of this proteostasis–respiration–redox axis sensitizes *C. jejuni* to both antibiotic and host-derived stress [14, 54, 55, 64, 74, 75]. More broadly, some antibiotic-tolerant states may depend less on passive shutdown than preservation of a minimal but critical homeostatic core under stress, particularly where proteostasis, respiration, and redox balance are tightly coupled. In this view, *C. jejuni* is informative because its reduced redundancy allows this ClpP-dependent homeostatic core to be resolved more clearly. ClpP-dependent proteostasis therefore emerges not simply as a species-specific determinant of tolerance, but as part of a broader mechanism of bacterial stress survival and a potentially useful antimicrobial target.

**Figure 6.**
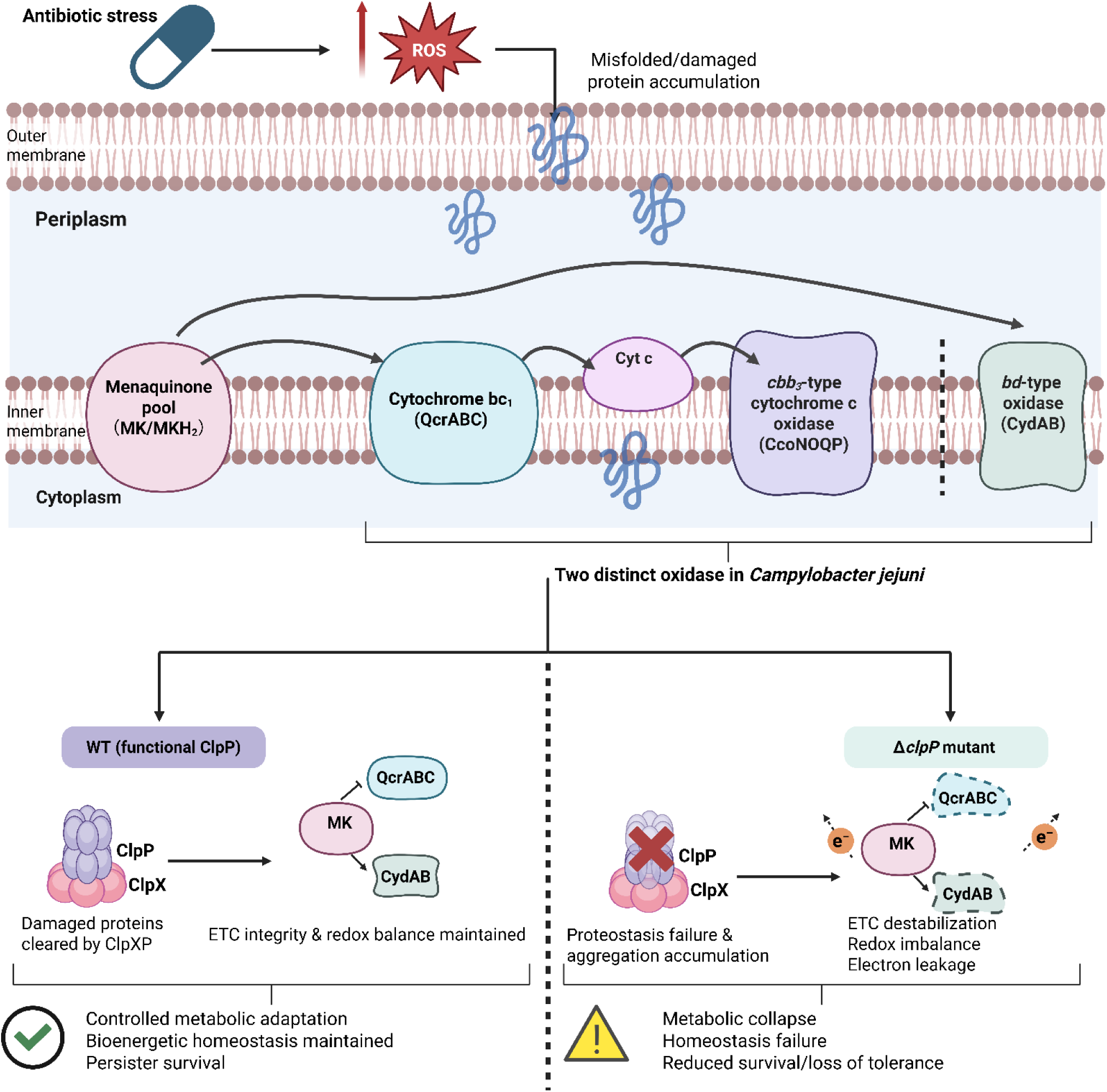
Proposed model of ClpP-dependent proteostasis and respiratory/redox homeostasis during antibiotic tolerance in *C. jejuni*. Under optimal microaerobic growth conditions, *C. jejuni* maintains efficient respiratory activity and energy conservation while ClpP-dependent proteostasis preserves protein integrity and supports overall cellular homeostasis. Upon exposure to bactericidal antibiotics and associated oxidative stress, cells enter a stress-adaptive tolerant state characterized by controlled metabolic downshift, respiratory remodeling, redox-oriented metabolic adaptation, and increased reliance on protein quality control. In this state, ClpP-mediated clearance of damaged or misfolded proteins is proposed to preserve respiratory chain integrity, maintain terminal oxidase function, and sustain survival-supportive homeostasis (**Bottom Left**). In the absence of ClpP, damaged proteins accumulate and proteostasis becomes disrupted, leading to impaired respiratory adaptation, amplified redox imbalance, membrane-associated damage, and reduced survival capacity under both antibiotic-treated and host-associated conditions (**Bottom Right**). This model summarizes the major conclusions derived from the multi-omics, respiratory, oxidative stress, macrophage infection, and mouse colonization analyses in this study.

## MATERIALS AND METHODS

### Bacterial strains and growth conditions

*C. jejuni* strain F38011 is a representative clinical isolate obtained from a fecal sample from a patient with bloody diarrhea, and was used in the current study [87]. Three additional *C. jejuni* strains 81116, 87-95, and 11168 were used in the antibiotic killing assay. Bacteria were routinely cultivated either on Mueller-Hinton agar (MHA) or in Mueller-Hinton broth (MHB) supplemented with selective antibiotics (chloramphenicol or kanamycin) at 37°C under microaerobic conditions.

### Construction of *C. jejuni* mutants and complementation

The mutants of *C. jejuni* F38011 were generated by replacing the genomic coding region of the targeted gene with a drug resistance gene using homologous recombination. The sequences of the primers are listed in **Table S1**. The *lon* deletion mutant was generated by homologous recombination and insertion of a chloramphenicol resistance cassette (Cm^R^). The upstream and downstream regions flanking *lon* (*lon^up^, lon^down^*) were PCR-amplified using the primer pairs OLU182/OLU183 and OLU186/OLU187, respectively. The Cm^R^ gene was amplified from the plasmid pACYC184 using the primer pairs OLU184/OLU185. The above three PCR fragments were purified using the Gel/PCR purification kit (Froggabio, CA) and inserted into EcoRI/XbaI-digested pUC19 using the NEBuilder® HiFi DNA Assembly Cloning Kit. The plasmid pUC19-*lon^up^-Cm^R^-lon^down^*was introduced into *C. jejuni* strain F38011 via natural transformation. Transformants were selected on MHA supplemented with 2 µg/mL Cm (MHBCm2) and streaked on the MHACm8 agar to confirm Cm resistance. Deletion of *lon* was confirmed by PCR.

The *clpP* deletion mutant of *C. jejuni* was generated using a similar protocol as described above. Briefly, upstream and downstream regions flanking *clpP* (*clpP^up^, clpP^down^*) were PCR-amplified using the primer pairs OUL196/OLU197 and OLU200/OLU201, respectively. The Cm^R^ gene was amplified from pACYC184 using the primer pair OLU198/OLU199. The engineered plasmid pUC19-*clpP^up^-Cm^R^-clpP^down^* was transformed into *C. jejuni* strain F38011, followed by selection on the MHACm2 and MHACm8 agar, subsequently. Deletion of *clpP* was confirmed by PCR.

The complementary plasmids were derived from pRY107 (Kan resistance) vector. The full *lon* gene, including the coding region and ∼500 bp upstream of the start codon, was PCR-amplified using primers OLU228/OLU229. The PCR products were ligated with EcoRI/XbaI-digested pRY107 using the NEBuilder® HiFi DNA Assembly Cloning Kit. The complementary plasmid pRY107-*lon* was transformed into *C. jejuni Δlon* strain and the complementary transformants were selected on the MHAKan20 agar. To construct the complementary plasmid of *clpP*, the full *clpP* gene, including the coding region and ∼400 bp upstream of the start codon, was PCR-amplified using the primers OLU230/231 and cloned into pRY107 subsequently. The complementary plasmid pRY107-*clpP* was transformed into *C. jejuni ΔclpP* strain and the complementary transformants were selected on the MHAKan20 agar.

The *spoT* mutant of *C. jejuni* was generated using the similar technique above. Briefly, upstream and downstream regions flanking *spoT* (*spoT^up^, spoT^down^*) were PCR-amplified using the primer pairs *spoT*-FF*/spoT*-FR and *sopT*-RF/*sopT*-RR, respectively. The Kan^R^ gene was amplified from pUC18K2 using the primer pair *kan*-F/*kan*-R. The engineered plasmid pUC19-*spoT^up^-Kan^R^-spoT^down^* was transformed into *C. jejuni* strain F38011, followed by selection on *kan* (50 μg/mL) agar, subsequently. Deletion of *spoT* was confirmed by PCR.

The *recA* deletion mutant of *C. jejuni* was generated using a similar protocol as described above. Briefly, upstream and downstream regions flanking *recA* (*recA^up^*, *recA^down^*) were PCR-amplified using the primer pairs *recA*-FF/*recA*-FR and *recA*-RF/*recA*-RR, respectively. The Cm^R^ gene was amplified from pRY111 using the primer pair *cm*-F/*cm*-R. The engineered plasmid pUC19- *recA^up^*-Cm^R^- *recA^down^* was transformed into *C. jejuni* strain F38011, followed by selection on the MHAkan50 agar subsequently. Deletion of *recA* was confirmed by PCR.

### Antibiotic killing and isolation of persister cells

Overnight *C. jejuni* culture was diluted to ∼log 8 CFU/mL in MHB containing either ampicillin (100 µg/mL) or ciprofloxacin (1 µg/mL). The bacterial culture was kept with shaking at 37℃ in a microaerobic condition. At indicated time points, aliquots were collected, centrifuged, washed with PBS, and plated on MHA. After incubation for 24 hr, 10-100 colonies were enumerated for each data point unless further noted. Along with antibiotic exposure, development of a biphasic killing curve could confirm the formation of *C. jejuni* persister cells.

### Determination of antibiotic MIC

MIC was determined using the broth microdilution method. Overnight bacterial culture was diluted to ∼log7 CFU/mL in MHB and added into 96-well flat-bottom clear polypropylene plates (Corning) with two-fold dilutions of antibiotic in a total volume of 100 µL. Plates were incubated at 37°C for 24 hr and then the absorbance at OD_600_ was measured using a Spectramax M3 microplate reader.

### Lipid peroxidation

Lipid peroxidation in *C. jejuni* cells was determined using the BODIPY™581/591 C11 sensor kit (Invitrogen™) according to the manufacturer’s instructions. Briefly, overnight cultures of wild-type *C. jejuni* F38011 and mutant strains were stained with dye in the dark for 1 hr and then added to 96-well black microplates (∼log 8 cells per well). Cells were treated with hydrogen peroxide (100 µM), ampicillin (100 µg/mL), or ciprofloxacin (1 µg/mL) for 2 hr. Lipid peroxidation level was evaluated as the relative increase in fluorescence emission at 525 nm.

### RNA-seq analysis

Total RNA of both untreated *C. jejuni* cells and persister cells was extracted using the RNeasy Mini Kit (Qiagen). Ribosomal RNA was removed by the RiboMinus Transcriptome Isolation Kit (Invitrogen) and mRNA was then purified using the RNeasy Mini Spin columns (Qiagen). The resulting transcriptome RNA was used for the construction of RNA-seq library using NEB Next mRNA Library Prep Reagent set from Illumina (NEB) according to the manufacturer’s protocol. These constructed libraries were sequenced by Illumina HiSeq 2000 platform by paired-end chemistry. Sequence reads processing, mapping to *C. jejuni* NCTC 11168 genome (NC_002163), and quantification of differential expression (RPKM) were conducted using CLC genomic workbench. The functional annotation of genes was conducted in the Kyoto Encyclopedia of Genes and Genomes (KEGG) database. Statistical significance is referred to as absolute log 2-fold change >1 and *p*-value <0.5.

### Quantitative PCR analysis

*C. jejuni* cells from both untreated cultures and cultures exposed to antibiotics for 4 hr were collected for RNA extraction using the RNeasy mini kit (Qiagen). Total RNA concentration and purity were determined using a Nanodrop 2000 Spectrophotometer (Thermo Scientific). The cDNA was produced via SuperScript™ II Reverse Transcriptase (Invitrogen). qPCR was performed using cDNA from bacterial samples with Power SYBR green PCR Master Mix (Applied Biosystems, Warrington, United Kingdom) on an ABI Prism 7000 Fast instrument (Life Technologies). The primers used for qPCR validation are listed in **Table S1**. Housekeeping genes *gyrA* and *rpoA* in *Campylobacter* were used as the internal controls. A total of 7 genes that were detected as differentially expressed in the RNA-seq results were selected for validation.

### Label-free quantitative proteomics by LC-MS/MS

*C. jejuni* persister cells were collected and cell pellets were suspended in ammonium bicarbonate (100 mM) and lysed by bead beating on ice. The lysates were diluted and digested with trypsin. Digested peptides were desalted, and their concentration was determined by the bicinchoninic acid protein assay. The sample was then analyzed by LC-MS/MS using a Waters nanoACQUITY UPLC® system (Milford, United States) coupled with a Q Exactive HF hybrid quadrupole-Orbitrap mass spectrometer (Thermo Scientific, San Jose, United States). Raw mass spectrometry data were processed using the MaxQuant proteomics platform [88]. The false discovery rate was set at 0.01 at the peptide and protein levels. Proteins were identified with at least 2 peptides of a minimum length of 6 amino acids in Uniprot database using *C. jejuni* NTCT 11168 proteome (UP000031762). Protein relative abundance was quantified using the MaxQuant-LFQ algorithm [89]. The sample with a measured LFQ intensity in at least two out of the three samples was subjected to quantification. Statistical significance is referred to as an absolute log 2-fold change>1 and a *p*-value for a two-tailed t-test <0.05.

### COG category annotation

PSI-BLAST (version 2.2.31+) was used to categorize proteins of *C. jejuni* F38011 according to the COG database with a default cut-off value [89]. Top PSI-BLAST hits were manually curated.

### Metabolomics and lipidomics

Both untreated and persister cells were subjected to dual extraction for both hydrophilic metabolomics and hydrophobic lipidomics. Briefly, 10 μL of ^13^C labeled hydrophilic metabolites and 10 μL of odd-chain lipid molecules were added to bacterial samples as the internal standards. Then, 270 μL of ice-cold methanol (kept at -20°C) was added to quench cell metabolism. The metabolites were released by three freeze-thaw cycles, including flash freezing in liquid nitrogen for 1 min and sonication in ice water for 15 min. After adding 900 μL of methyl tert-butyl ether (MTBE), the samples were sonicated in ice water for 30 min to extract hydrophobic molecules. Next, the samples were mixed with 315 μL of cold H_2_O and centrifuged at 20,817 ×*g* and 10°C for 2 min to achieve a clear phase separation between the hydrophilic methanol-H_2_O layer (bottom) and the hydrophobic MTBE layer (top). The MTBE layer was collected, speed vacuum dried, reconstituted with 100 μL of isopropanol alcohol/acetonitrile (1:1, v/v), and stored at - 80°C for further lipidomic analysis. The methanol-H_2_O layer of the samples was kept at -20°C for 2 hr to precipitate the proteins. After centrifugation at 20,817 ×*g* and 4°C for 15 min, the supernatant was collected. This aqueous solution was dried in a speed-vacuum drier, reconstituted using 100 μL of acetonitrile/H_2_O (1:1, v/v), and stored at -80°C for further metabolomic analysis. Quality control samples for both metabolomics and lipidomics were prepared by mixing 10 μL of each sample. One method-blank sample (*i.e.*, an empty tube subjected to the overall extraction procedure) and two solvent-blank samples (*i.e.*, reconstitution solvent) were prepared as the negative controls.

Metabolomic (positive and negative) and lipidomic (negative) analyses of the samples were conducted using an Agilent 1290 infinite UHPLC system (Agilent Technologies, Palo Alto, CA) coupled with a Bruker Impact II quadrupole-time-of-flight (QTOF) mass spectrometer (Bruker Daltonics, Bremen, Germany). For hydrophilic metabolomic analysis, samples were separated on a SeQuant ZIC-pHILIC column (2.1 × 150 mm, 3.5 μm, 100 Å). Mobile phase A contained 5 mM NH_4_Ac in 95% H_2_O and 5% acetonitrile and mobile phase B constituted of 5 mM NH_4_Ac in 5% H_2_O and 95% acetonitrile. The pH of mobile phase A was adjusted to 5 for positive mass spectral collection, and the pH was maintained at 9.8 for negative mass spectral collection. Sample elution was achieved using a linear gradient at 30°C with a flow rate of 200 μL/min. Specifically, the gradient started with 95% B for the first 3 min, decreased to 75% B over 4 min, then to 25% B at 12 min, again to 5% B at 14 min, and finally maintained at 5% for 4 min. The column was then equilibrated at 95% B for 18 min before the injection of the next sample. Sample injection volume was optimized using a quality control sample to maximize spectral intensity and avoid signal saturation.

For lipidomic negative analysis, samples were separated on a Waters Acquity UPLC BEH C_18_ column (1 × 100 mm, 1.7 μm, 130 Å). The mobile phases for the separation were A: 5 mM ammonium formate in acetonitrile/H_2_O (6:4 v/v) and B: 5 mM ammonium formate in isopropanol alcohol/acetonitrile (9:1, v/v). The gradient sample elution program started with 5% B, increased to 40% B at 8 min, again to 70% B at 14 min, finally reached 95% B at 20 min, and maintained for 3 min. The column was then equilibrated at 5% B for 9 min before the next injection. The column was maintained at 25°C with a flow rate of 150 μL/min. Sample injection volume was optimized using the same approach as that for metabolomic analysis. An electrospray ionization source was used to ionize metabolites for MS detection and spectral information was recorded using the data-dependent MS/MS approach (auto MS/MS). The spectra for auto MS/MS were collected using a spectral rate of 8 Hz and *m/z* range of 50 to 1,200. For the auto MS/MS method, the MS/MS spectra were acquired using a stepping collision energy of 40 eV to 120 eV, an intensity threshold of 300, a cycle time of 3 s, a smart exclusion after two spectra, and a precursor reconsideration for intensity difference >1.8. The data- dependent LC-MS/MS results were normalized to total protein content according to the Bradford assay and processed using MS-DIAL (RIKEN, version 3.52). All the metabolic features with MS/MS spectra were manually confirmed using the MS-Finder (RIKEN, version 3.20), METLIN (Scripps Research), or an in-house metabolic library with an m/z tolerance of 5 ppm. Significantly regulated metabolic features were defined as absolute log 2-fold >2 and *p*-value for two-tailed t-test <0.05.

### Gene set enrichment analysis

Gene set enrichment analysis (GSEA) was conducted using the Java (GSEA 3.0)-based graphical user interface with a pre-ranked list of the classic enrichment statistics [90, 91]. GSEA was used to determine whether a pre-defined set of genes/proteins is significantly different between persister cells and untreated cells. The entire list of genes/proteins was ranked according to the expression difference between persister cells and untreated cells. The cumulative distribution function was constructed by performing 1000 random gene set membership assignments.

### Visualization of the enrichment map

The significantly enriched terms in transcriptome and proteome analysis were visualized in Cytoscape 3.7.1 [92]. Enrichment maps were built with stringent pathway similarity scores (Jaccard and overlap combined coefficient 0.6) and manually curated for the most representative groups of similar pathways and biological processes.

### Seahorse extracellular flux analysis

Extracellular flux analysis was performed using an XFe96 Extracellular Flux Analyzer to measure bacterial respiration as oxygen consumption rate (OCR). This assay was adapted from a previously published method with modifications [34]. Overnight cultures of *C. jejuni* F38011 wild-type and the *ΔclpP* were grown in MHB at 37°C with shaking at 175 rpm in a CO_2_ incubator shaker supplied with 10% CO_2_. Cultures were grown to an OD_600_ of ∼0.3 and then diluted in fresh MHB to an OD_600_ of 0.06, corresponding to ∼2 × 10^8^ CFU/mL. For plate preparation, XF cell culture microplates were coated with poly-D-lysine (PDL). Briefly, 15 μL of 1 mg/mL PDL prepared in 100 mM Tris-HCl (pH 8.4) was added to each well, followed by overnight drying in a biosafety cabinet and two rinses with distilled water. Then, 90 μL of diluted bacterial suspension was added to each well. Cells were attached to the PDL-coated wells by centrifugation at 3,313 x *g* for 10 min in a benchtop swinging-bucket centrifuge to generate a monolayer of adhered cells. After centrifugation, 90 μL of MHB was gently added to each well to bring the final pre-injection volume to 180 μL. Following calibration with equilibration enabled, basal OCR was recorded before antibiotic addition to confirm uniform cell seeding. The baseline phase consisted of 3 measurement cycles, each including 1.5 min of mixing, 1.5 min of waiting, and 3 min of measurement for a total baseline duration of 18 min. For treatment conditions, 20 μL of 10× antibiotic stock was injected through port A to achieve final concentrations of 20 μg/mL ampicillin or 0.2 μg/mL ciprofloxacin. For untreated controls, 20 μL of MHB alone was injected. OCR was then monitored using 30 measurement cycles, each consisting of 1.5 min of mixing, 1.5 min of waiting, and 3 min of measurement for a total post-injection duration of 3 hr. Wells receiving MHB alone served as no-treatment controls.

OCR values were normalized to the number of viable cells, expressed as CFU/mL, determined by plating assay. For normalization, bacteria were cultured and treated in parallel on an XF Cell Culture Microplate under the same conditions as those used for the Seahorse assay. At 30 min after the aerobic incubation, aliquots were collected from parallel wells, serially diluted in sterile PBS, and plated onto MHA for colony enumeration. After incubation under microaerophilic conditions at 37°C for 48 hours, colonies were counted and viable cell numbers were calculated as CFU/mL. OCR values were then normalized to the corresponding viable cell counts obtained from matched treatment groups.

### Protein extraction and Western blot analysis

*C. jejuni* cells grown in MHB were harvested by centrifugation, and total protein was extracted using B-PER II bacterial protein extraction reagent supplemented with EDTA-free protease inhibitor, lysozyme, and DNase I according to the manufacturer’s instructions. Protein samples were prepared in LDS sample buffer supplemented with reducing agent and separated on 4-12% Bis-Tris gels in MES SDS running buffer with antioxidant added to the inner chamber at 120 V for 100 min. Proteins were then transferred onto 0.45-μm PVDF membranes. Before transfer, membranes were activated in methanol for 1 min, and gels were equilibrated in Towbin transfer buffer without SDS or methanol for 10 min. Transfer was performed using a Trans-Blot Turbo Transfer System in Towbin transfer buffer without SDS and methanol. Membranes were blocked in 5% bovine serum albumin in PBST for 1 hr at room temperature and incubated overnight at 4°C with anti-CydA antibody diluted 1:5,000 in 5% bovine serum albumin in PBST. After three washes with PBST for 10 min each, membranes were incubated for 1 hr at room temperature with HRP-conjugated goat anti-rabbit secondary antibody diluted 1:5,000 in 5% bovine serum albumin in PBST, followed by five washes with PBST for 10 min each. Protein bands were detected using enhanced chemiluminescence substrate. Membranes were incubated with substrate for 1 min in the dark and imaged by chemiluminescence. Ponceau S and chemiluminescence images were merged for figure preparation.

### RAW 264.7 macrophage infection assay

RAW 264.7 murine macrophages were maintained in Dulbecco’s modified Eagle medium supplemented with 10% fetal bovine serum at 37°C in a humidified incubator with 5% CO_2_ as previously described with modifications. For infection assays, cells were seeded in 24-well plates and infected with *C. jejuni* F38011 wild-type or the *ΔclpP* mutant at a multiplicity of infection (MOI) of 100 [93]. Following infection, macrophages were incubated for 3 hr to assess bacterial adherence. At the indicated time points, monolayers were washed twice with PBS to remove non-adherent bacteria. For adherence assays, macrophages were lysed with 0.1% Triton X-100, and recovered bacteria were serially diluted and plated for CFU enumeration. To assess bacterial survival in the host-associated environment, infected macrophages were further incubated until the indicated time points, lysed, and viable bacteria were quantified by the plating assay. Bacterial counts were expressed as CFU per well and compared between strains.

### Mouse colonization assay

To determine the contribution of ClpP to host colonization, 8-week-old Sigirr^-/- mice were orally inoculated with *C. jejuni* 81-176, F38011 wild-type or the *Δlon* and *ΔclpP* mutants at a dose of ∼1 × 10^7^ CFU per mouse, following an established protocol [52]. At the indicated timepoints post-infection, mice were euthanized, and intestinal tissues and/or intestinal contents were collected. Samples were suspended in sterile PBS, homogenized, serially diluted, and plated on MHA containing selective supplements. Plates were incubated for 48 hr at 42°C under microaerobic conditions, after which colonies were enumerated. Bacterial burdens were calculated and expressed as CFU per gram of tissue or intestinal content. Animal experiments were conducted under protocol A11-290, approved by the University of British Columbia Animal Care Committee, in accordance with Canadian Council of Animal Care (CCAC) guidelines. Mice were monitored throughout infection and were euthanized upon signs of severe distress or body weight loss exceeding 15%.

### Histology, pathological scoring and immunofluorescent staining

Histological analysis was conducted according to an established protocol with minor modifications [52]. Intestinal tissues were collected from infected mice, fixed in 10% neutral-buffered formalin, paraffin embedded, and sectioned. For histological evaluation, tissue sections were stained using hematoxylin and eosin and examined microscopically for the presence and localization of *C. jejuni* at the intestinal surface.

## Supporting information

Correlation analysis

Disorder analysis

Lipidomics

Metabolomics

RNA seq

## ACKNOWLEDGEMENTS

This work was supported by funds provided to X.L. from Natural Sciences and Engineering Research Council of Canada (NSERC RGPIN-2019-03960, RGPIN-2019-00024, RGPIN-2025- 03996), Canada Research Chairs Program (CRC-2024-00011), Sector Innovation Program of Genome Canada and Genome British Columbia, Genomics Integration Program of Genome Québec, Fonds de recherche du Québec – Nature et technologies, and Canadian Poultry Research Council.

## EXTENDED DATA

**Extended Data Figure 1.**
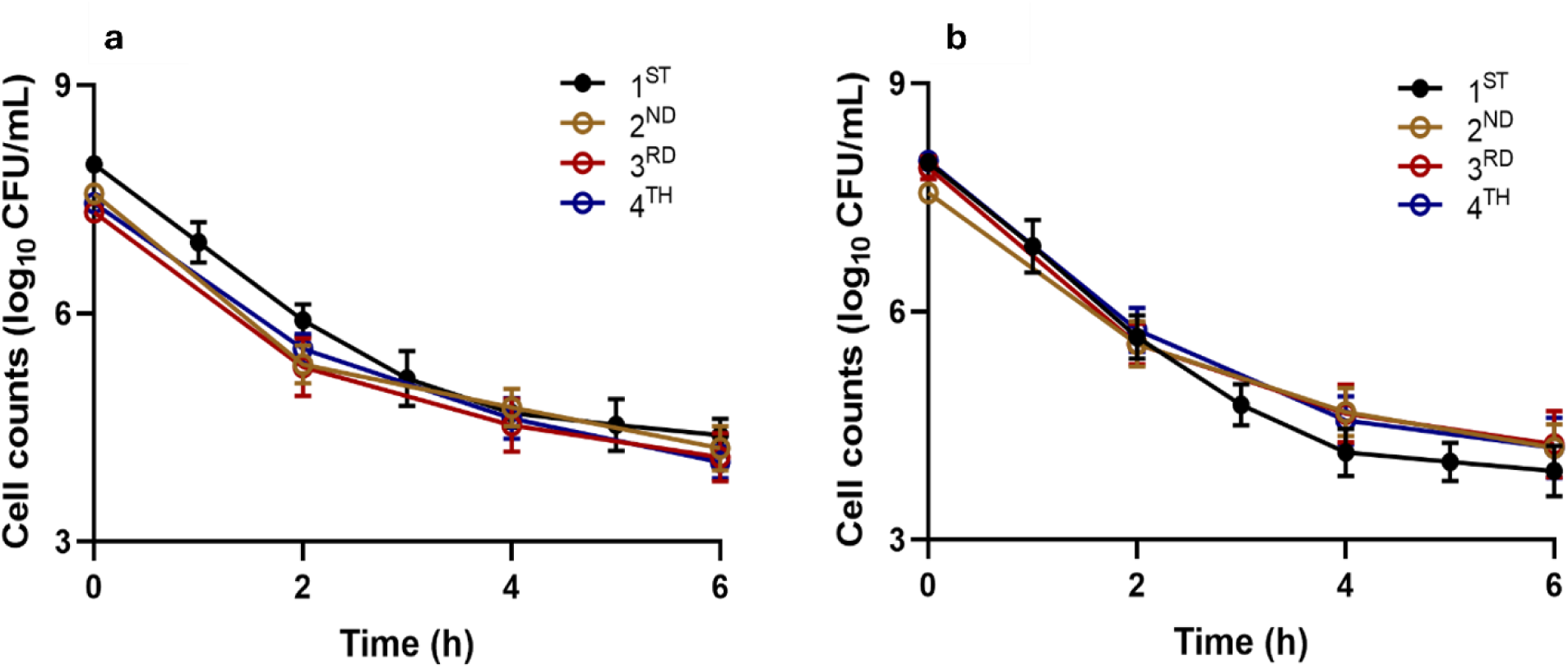
*C. jejuni* F38011 wild type strain were subjected to multiple rounds of challenges with ampicillin (a) and ciprofloxacin (b) for multiple rounds, and no detectable resistance observed.

**Extended Data Figure 2.**
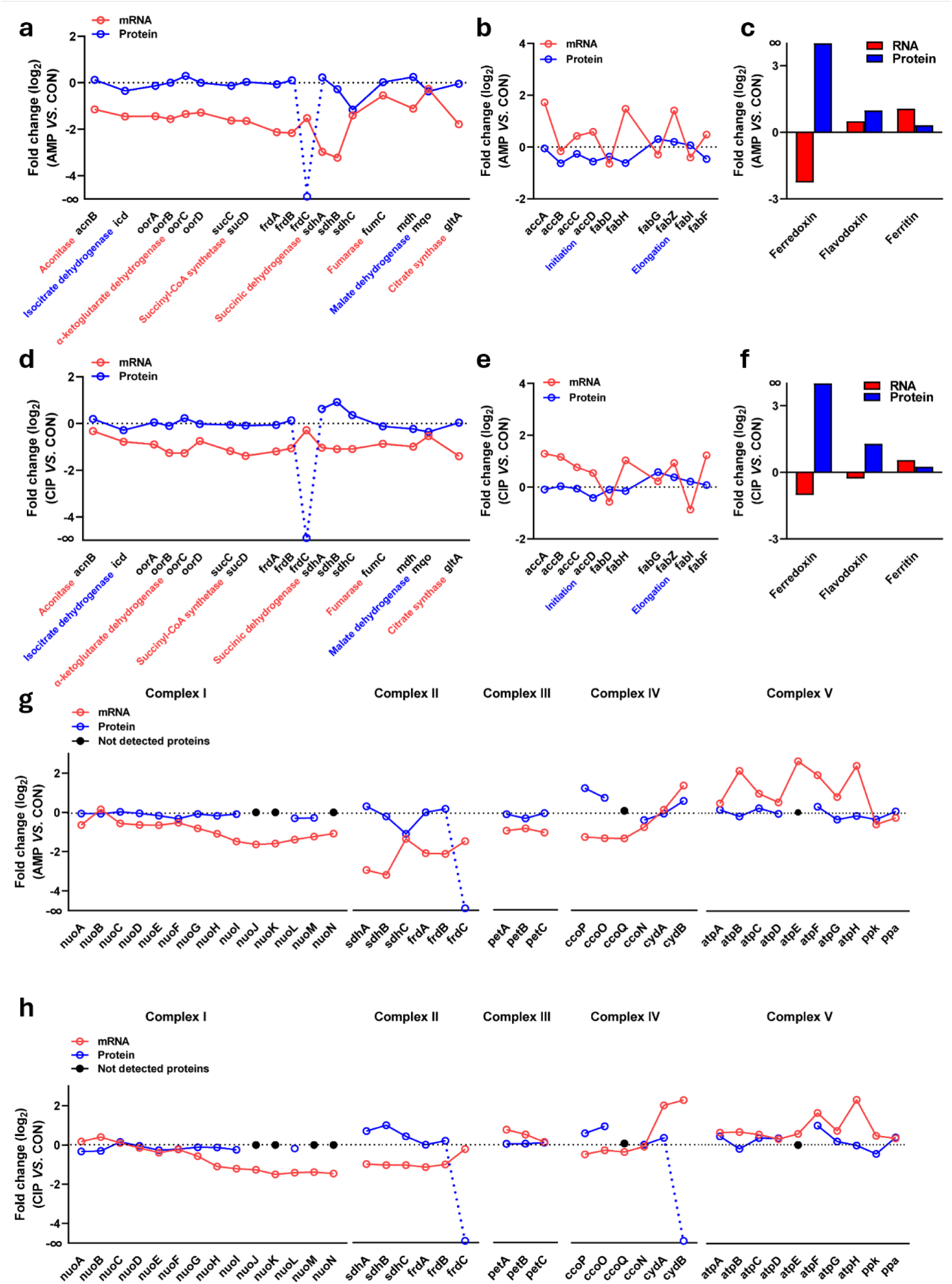
Transcriptomic and proteomic remodeling of central metabolism, fatty acid biosynthesis, redox-associated factors and respiratory complexes in antibiotic-tolerant *C. jejuni*. (a, b) Log2 fold changes in mRNA and protein abundance of selected tricarboxylic acid (TCA) cycle and associated metabolic enzymes in ampicillin (AMP)-selected (a) and ciprofloxacin (CIP)-selected (b) tolerant cells relative to untreated control cells (CON). (c, d) Log2 fold changes in mRNA and protein abundance of selected fatty acid biosynthesis enzymes in AMP-selected (c) and CIP-selected (d) tolerant cells relative to untreated controls. (e, f) Log2 fold changes in mRNA and protein abundance of selected redox- and iron-associated factors, including ferredoxin, flavodoxin and ferritin, in AMP-selected (e) and CIP-selected (f) tolerant cells relative to untreated controls. (g, h) Log2 fold changes in mRNA and protein abundance of subunits of respiratory complexes I–V in AMP-selected (g) and CIP-selected (h) tolerant cells relative to untreated controls. Red, mRNA; blue, protein. Black dots indicate proteins not detected in the proteomic dataset. Features detected exclusively by transcriptomics, with no corresponding detection in the proteomics, are annotated as -∞ fold change for visualization purposes.

