## Supplemental figure 1_7, and tables 1_2 for "ClpP-dependent protein quality control supports antibiotic persistence in *Campylobacter jejuni* through bioenergetic homeostasis"

**Supplementary Information**

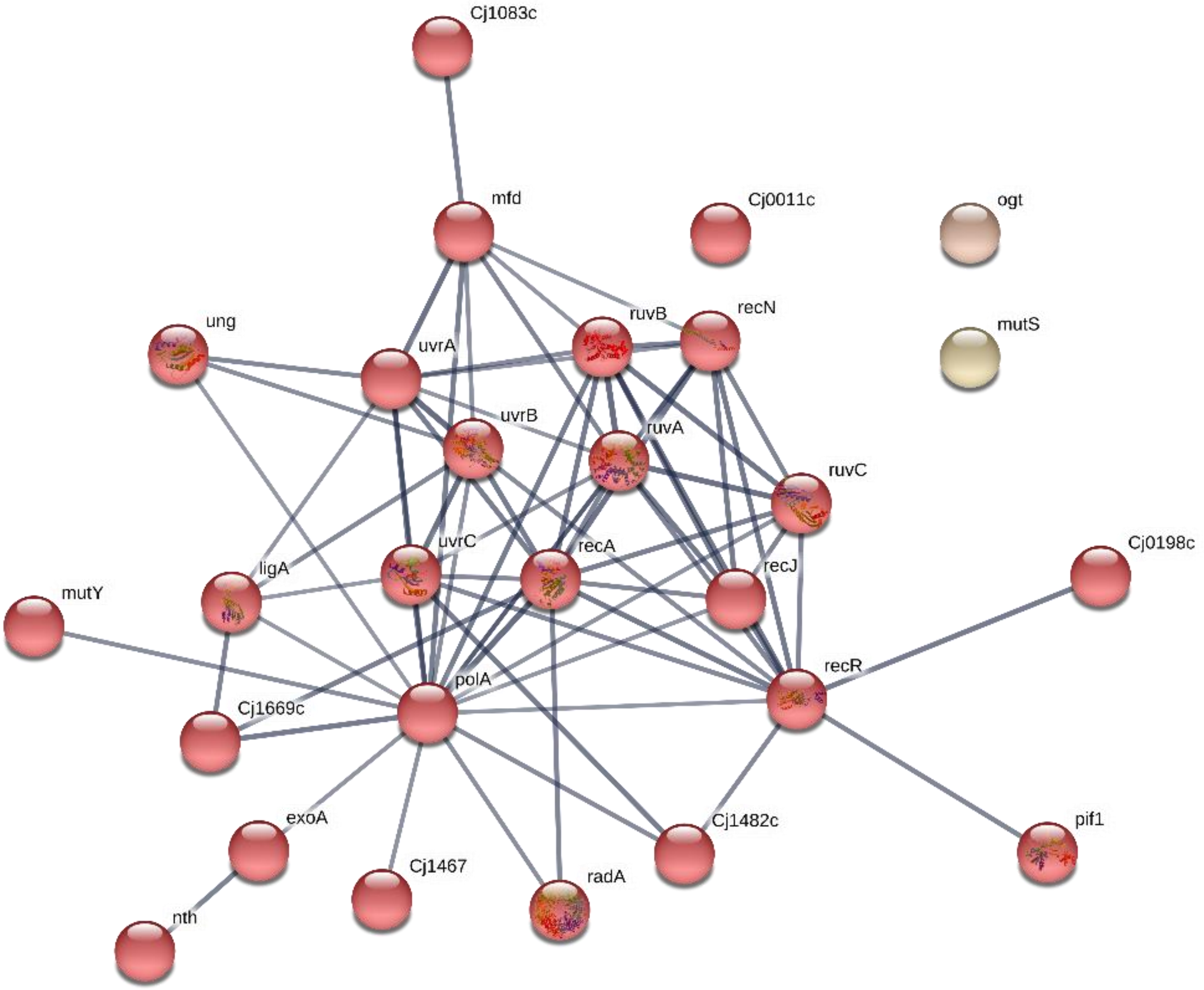


**Figure S1. Network analysis was conducted to predict gene interactions within the DNA repair pathway.** The selected genes were the major components of DNA repair GO terms GO:0006281 and GO:0009432. The network was built using the STRING database. The genes were clustered in k-means. Colored edges represented the type of associations as indicated in the STRING database.

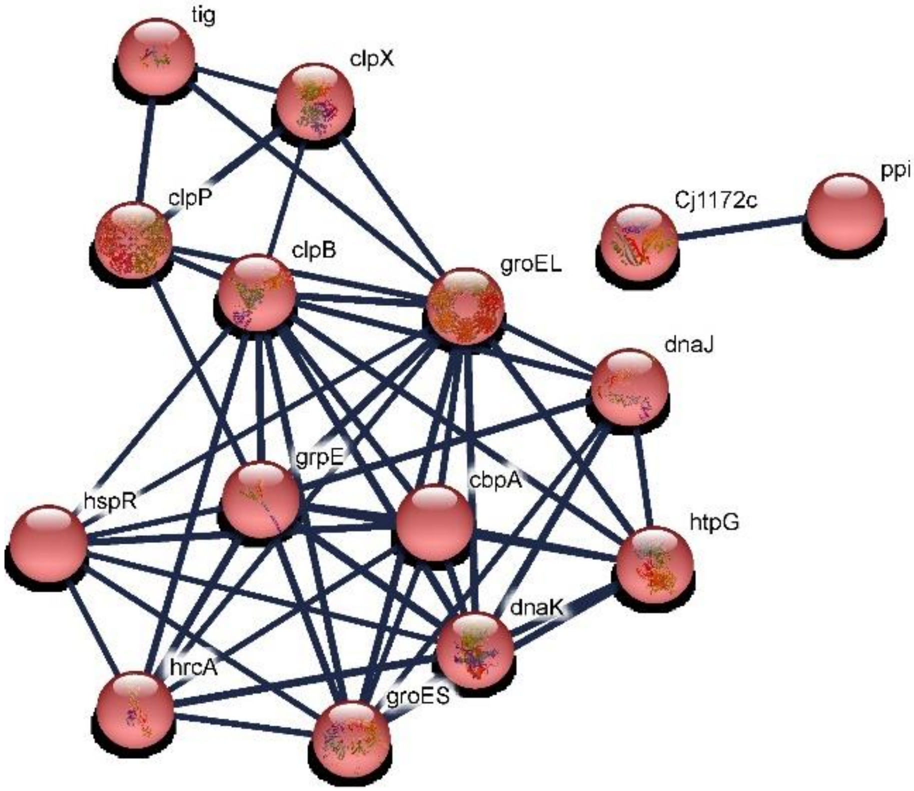


**Figure S2. Network analysis was conducted to predict gene interactions within the protein folding pathway.** The selected genes were the major components of DNA repair GO terms GO:0006457. The network was built using the STRING database. The genes were clustered in k-means. Colored edges represented the type of associations as indicated in the STRING database.

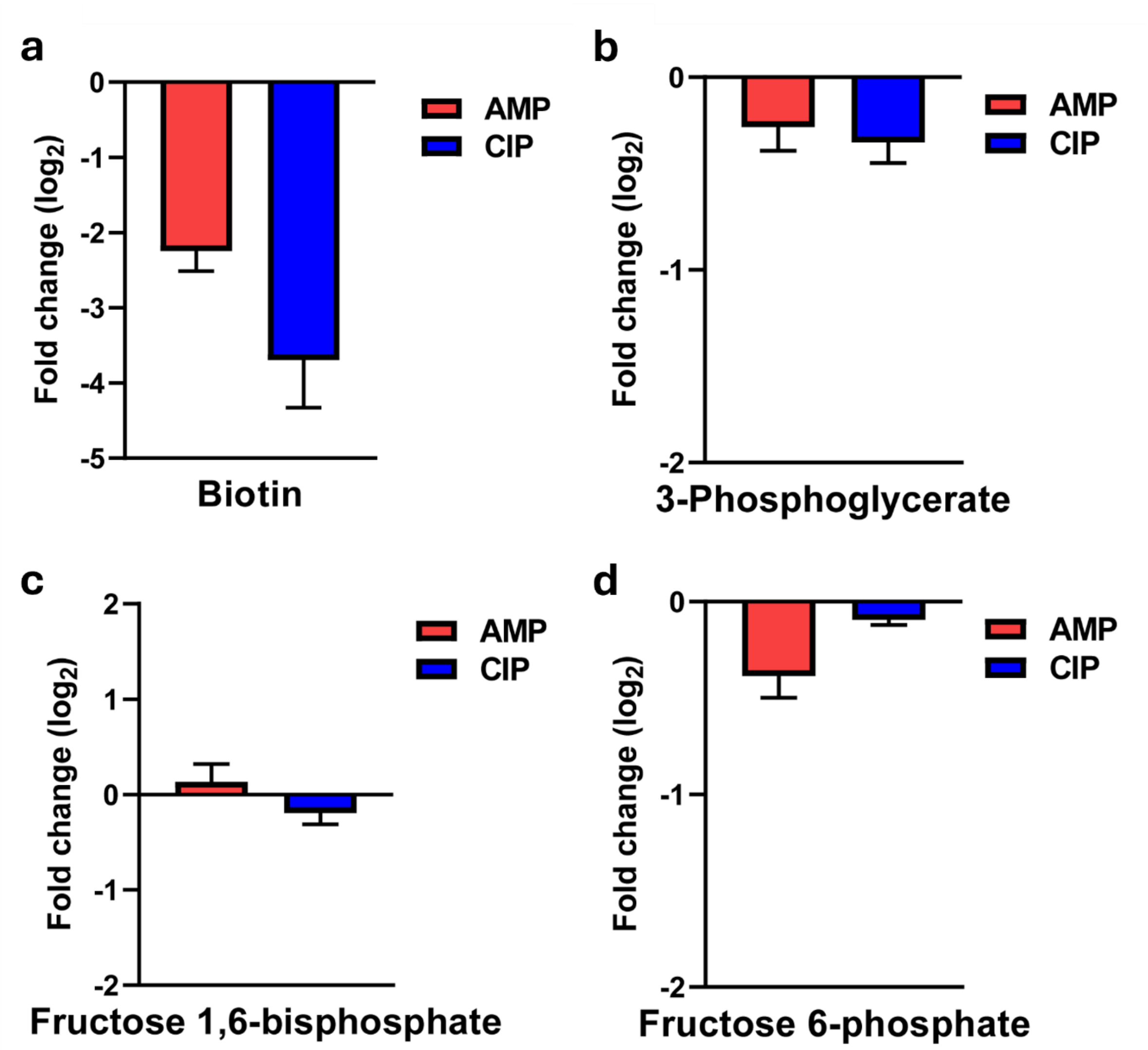


**Figure S3. Metabolic analysis of *C. jejuni* F38011 persisters showed distinct metabolomic profile.** Biotin synthesis (A), metabolic intermediate in glycolysis: 3-phosphoglyceric acid (B), fructose 1,6-bisphosphate (C), and fructose 6-phosphate (D).

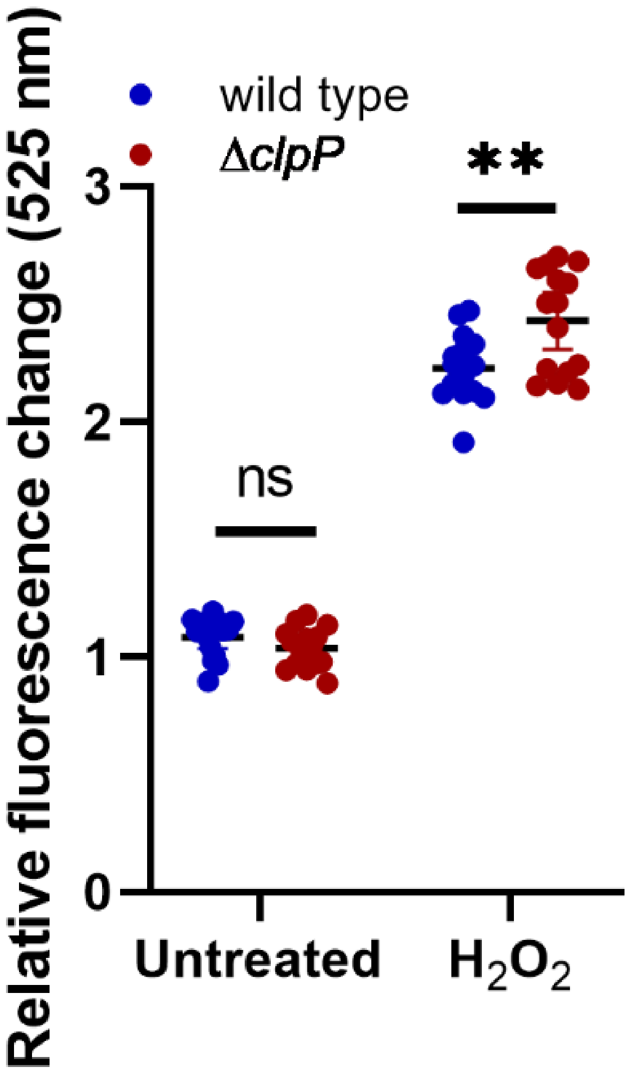


**Figure S4. Lipid peroxidation of wild-type and ΔclpP mutant *C. jejuni* cells was determined using the BODIPY™581/591 $C_{11}$ sensor kit (Invitrogen™).** *C. jejuni* wild-type and Δ*clpP* mutant showed significant differences in lipid oxidation after hydrogen peroxide treatment (100 µM).

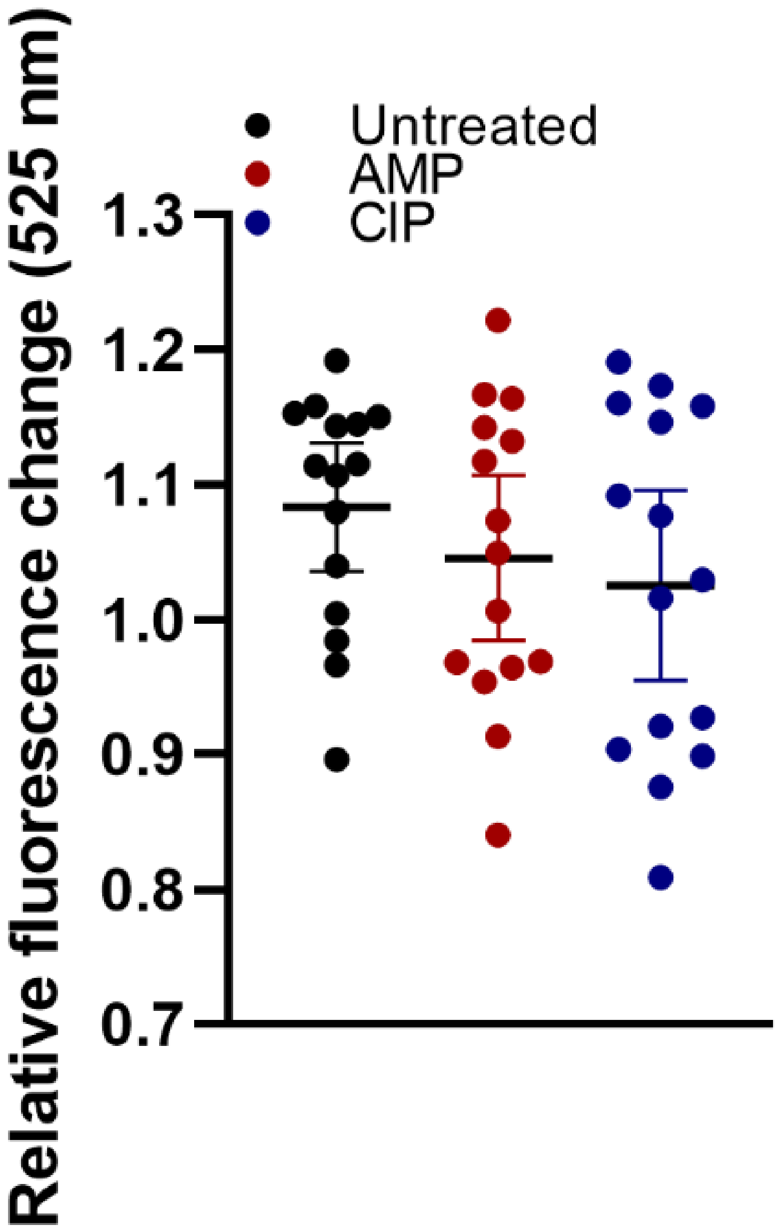


**Figure S5. Lipid peroxidation of *C. jejuni* persister cells.** F38011 exposed to either ampicillin or ciprofloxacin did not reveal significant difference in lipid peroxidation.

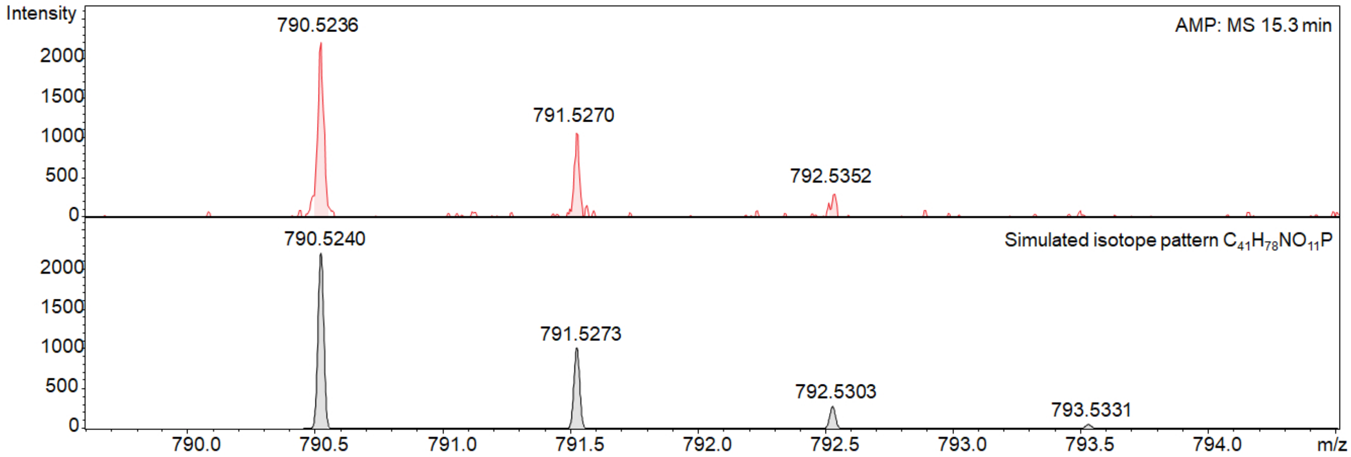


**Figure S6. Comparison of the theoretical isotope pattern calculation of $C_{41}H_{78}NO_{11}P$ (bottom) and one oxidized phosphatidylethanolamine observed experimentally (top) in ampicillin treated *C. jejuni* cells.**

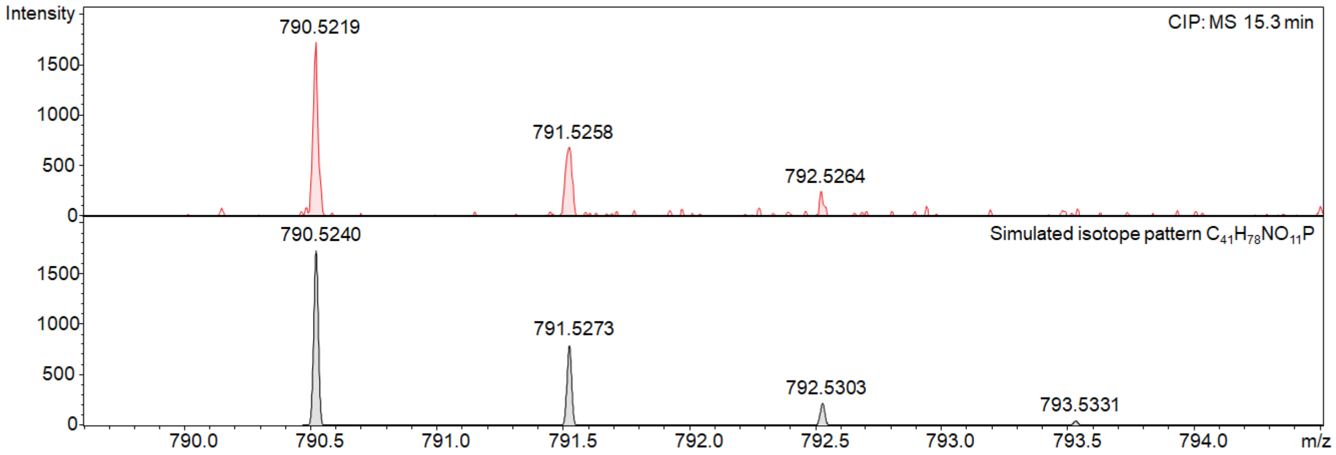


**Figure S7. Comparison of the theoretical isotope pattern calculation of $C_{41}H_{78}NO_{11}P$ (bottom) and one oxidized phosphatidylethanolamine observed experimentally (top) in ciprofloxacin treated *C. jejuni* cells.**

**Table S1. Bacterial strains and measured MIC**

| Strains | Description | Reference | MIC (µg/ml) | |
|---|---|---|---|---|
| | | | Ampicillin | Ciprofloxacin |
| *C. jejuni* 81116 | Human clinical isolate | Neal-McKinney et al., 2010 [1] | 8 | 0.08 |
| *C. jejuni* 87-95 | Laboratory strain from the Dr. M. Blaser collection (Vanderbilt University) | Konkel et al., 1997 [2] | 8 | 0.08 |
| *C. jejuni* 11168 | Human clinical isolate | Parkhill et al., 2000 [3] | 8 | 0.08 |
| *C. jejuni* F38011 | Human clinical isolate | Feng et al., 2016 [4] | 8 | 0.08 |
| *C. jejuni* F38011 Δ*clpP* | mutant | This study | 8 | 0.08 |
| *C. jejuni* F38011 Δ*clpP*:: *clpP* | mutant | This study | 8 | 0.08 |
| *C. jejuni* F38011 Δ*Lon* | mutant | This study | ND* | ND* |
| *C. jejuni* F38011 Δ*recA* | mutant | This study | ND* | ND* |
| *C. jejuni* F38011 Δ*spoT* | mutant | This study | ND* | ND* |

*ND, not determined. MICs for the Δ*lon*, Δ*recA*, and Δ*spoT* mutants were not determined because these mutants did not show altered persistence phenotypes relative to the parental strain in preliminary assays under the conditions tested and were therefore not further characterized.

**Table S2. Primers used for cloning.**

| Primer sets | Forward primer (5'---3’) | Reverse primer (5’---3’) | Explanation |
|---|---|---|---|
| OLU182 OLU183 | ttgtaaaacgacggccagtgaattcCGCAGAAGTTGCAAACCAATAC | tactgcgatgagtggcagggcggggcgtaaGGTACCGGGTGATGTAATGAAAGAATCAGCT | Upstream and downstream regions flanking lon ($lon^{up}$, $lon^{down}$). |
| OLU186 OLU187 | ttagcttccttagctcctgaaaatctcgatGGATCCCAAATCACAAATCCTAGACGCATC | caagcttgcatgcctgcaggtcgactctagaCAATCTTAGTGCTAGTGAGTGG | |
| OLU184 OLU185 | AGCTGATTCTTTCATTACATCACCCGGTACCttacgccccgccctgccactcatcgcagta | GATGCGTCTAGGATTTGTGATTTGGGATCCatcgagattttcaggagctaaggaagctaa | Primers to amplify chloramphenicol resistance gene ($Cm^R$) on the plasmid pACYC184 for *lon* mutant. |
| OUL196 OLU197 | ttgtaaaacgacggccagtgaattcGGGATGCCTGAAGGACTAAG | tactgcgatgagtggcagggcggggcgtaaGGTACCGCACAAGAAGCTAAAGAATATG | Upstream and downstream regions flanking *clpP* ($clpP^{up}$, $clpP^{down}$) |
| OLU200 OLU201 | ttagcttccttagctcctgaaaatctcgatGGATCCGCGTGAATAAATGTCATAAC | caagcttgcatgcctgcaggtcgactctagaCCAGGTTTTGAAGATGGTA | |
| OLU198 OLU199 | CATATTCTTTAGCTTCTTGTGCGGTACCttacgccccgccctgccactcatcgcagta | GTTATGACATTTATTCACGCGGATCCatcgagattttcaggagctaaggaagctaa | Primers to amplify chloramphenicol resistance gene ($Cm^R$) on the plasmid pACYC184 for *clp* mutant. |
| OLU228 OLU229 | ttgtaaaacgacggccagtgaattcGCGCAAGACATGTAGCCCTT | caagcttgcatgcctgcaggtcgactctagaGAGCATGTTGCCGATCCTTT | Coding region and ~500 bp upstream of the start codon of *lon* |
| OLU230 OLU231 | caagcttgcatgcctgcaggtcgactctagaTTTGATGCATTTGCAGTGCC | ttgtaaaacgacggccagtgaattcGTCTAGGATTGGGATAGGTGATAAT | Coding region and ~400 bp upstream of the start codon of *clp* |
| *spoT*-FF *spoT*-FR | CGGAATTCAAGTGGAGAGCCTTATGCGG | CTTGGTACCGTCTATGGGCTATTGGGGCA | Upstream and downstream regions flanking *spoT* ($spoT^{up}$, $spoT^{down}$) |
| *sopT*-RF *sopT*-RR | GCGGATCCAGCCAGACGTATTAGACAAGTAGC | GATCTAGATCTCAAAATAATCTACCGCCGA | |
